# Deep-learning based Embedding of Functional Connectivity Profiles for Precision Functional Mapping

**DOI:** 10.1101/2025.01.29.635570

**Authors:** Jiaxin Cindy Tu, Jung-Hoon Kim, Patrick Luckett, Babatunde Adeyemo, Joshua S. Shimony, Jed T. Elison, Adam T. Eggebrecht, Muriah D. Wheelock

## Abstract

Spatial correlation of functional connectivity profiles across matching anatomical locations in individuals is often calculated to delineate individual differences in functional networks. Likewise, spatial correlation is assessed across average functional connectivity profiles of groups to evaluate the maturity of functional networks during development. Despite its widespread use, spatial correlation is limited to comparing two samples at a time. In this study, we employed a variational autoencoder to embed functional connectivity profiles from various anatomical locations, individuals, and group averages for simultaneous comparison. We demonstrate that our variational autoencoder, with pre-trained weights, can project new functional connectivity profiles from the vertex space to a latent space with as few as two dimensions, yet still retain meaningful global and local structures in the data. Functional connectivity profiles from various functional networks occupy distinct compartments of the latent space. Moreover, the variability of functional connectivity profiles from the same anatomical location is readily captured in the latent space. We believe that this approach could be useful for visualization and exploratory analyses in precision functional mapping.

## 1 Introduction

Distributed large-scale networks in the human neocortex (Damoiseaux et al., 2006; M. D. Fox et al., 2005; Seitzman, Snyder, et al., 2019) with trait-like inter-individual variation in network topography studied extensively with resting-state functional MRI (fMRI) functional connectivity (FC)(Bijsterbosch et al., 2018; Braga & Buckner, 2017; Cui et al., 2020; Dworetsky et al., 2021, 2024; Glasser et al., 2016; Gordon, Laumann, Adeyemo, & Petersen, 2017; Gordon, Laumann, Adeyemo, Gilmore, et al., 2017; Gordon, Laumann, Gilmore, et al., 2017; Gratton et al., 2018; Kong et al., 2019; Kraus et al., 2021; H. Li et al., 2017; Seitzman, Gratton, et al., 2019; D. Wang et al., 2015). These functional networks have been attributed to distinct functional roles based on their strong spatial correspondence to the specialized functional systems activated during task fMRI (Cole et al., 2016; P. T. Fox & Friston, 2012; Power et al., 2011; Wig, 2017; Yeo et al., 2011). Due to the close relationship between the resting-state networks and functional significance, precision functional mapping to capture individual-specific functional networks demonstrates the potential to advance both psychiatric research and personalized therapeutic interventions (M. D. Fox et al., 2013; Gratton, Kraus, et al., 2020; Labonte et al., 2024; Lynch et al., 2022, 2024).

Many precision functional mapping methods involve the comparison of FC profiles from individual seed locations (e.g. vertices or voxels) in different brains. Reliable mapping of individual-specific functional networks requires a long fMRI data acquisition(Gordon, Laumann, Gilmore, et al., 2017). Therefore, many researchers have opted to use a group consensus network as a prior and compare the FC profiles to generate individualized functional networks (Gordon, Laumann, Adeyemo, & Petersen, 2017; Kong et al., 2019; H. Li et al., 2017). For example, one technique called “template matching” (Gordon, Laumann, Adeyemo, & Petersen, 2017; Hermosillo et al., 2024; Moore et al., 2024) assigns network identities to individual locations based on the similarity of the best matching network average FC profiles. Another method identifies trait-like “network variants” as contiguous cortical regions with low spatial correlation between an individual FC profile and a group average FC profile from the anatomically matched seed locations (Seitzman, Gratton, et al., 2019). In a closely related school of analyses, seed-based FC profiles were compared across group-average FC data to measure FC “maturity” (i.e. similarity to adults) in pediatric cohorts (Gao et al., 2015; Sylvester et al., 2022). Despite the prevalence of using a scalar summary of the similarity between FC profiles, this does not demonstrate the specific connections that drive the similarity/dissimilarity, nor whether the number and topography of networks in the original set of network priors are appropriate. Moreover, the “network variant” and “maturity” measures only compare FC profiles with direct anatomical correspondence, yet a profound literature of evidence suggests that functional correspondence across subjects may not perfectly align with anatomical correspondence (Guntupalli et al., 2018; Haxby et al., 2020).

Dimensionality reduction can be beneficial for efficiently comparing multiple samples of high-dimensional data, such as FC profiles. Various dimensionality reduction methods have been applied in neuroscience research to visualize and gain insight from high-dimensional data, including animal behavior (Stringer et al., 2019), RNA-sequencing (Zeisel et al., 2015), neural population activity (Churchland et al., 2012; Pandarinath et al., 2018), fMRI activity (Calhoun et al., 2001; Gotts et al., 2020; Kriegeskorte et al., 2008; Pospelov et al., 2021; Smith et al., 2004), and fMRI functional connectivity (Hacker et al., 2013; Margulies et al., 2016). In particular, the functional gradient calculated using diffusion map embedding on functional connectivity data has provided insights into the relative similarity of FC profiles from all seed locations in one individual or group-averaged brain in many recent studies (Dong et al., 2021; Hong et al., 2019; Langs et al., 2010, 2016; Larivière et al., 2020; Margulies et al., 2016; Nguyen et al., 2023; Tian et al., 2020; Xia et al., 2022). While most dimensionality reduction methods can effectively represent the relationship between the existing data samples in a low-dimensional latent space, few can back project the data from the low-dimensional latent space easily to the original data space, especially for new data samples outside the original data distribution. Moreover, existing methods to compare across individuals were based on first finding a low-dimensional latent space for each individual, and then applying Procrustes analysis (Bookstein, 1997) to align the spaces across subjects such that “the distance between a randomly chosen subset of vertices from the same anatomical part of the brain is minimized in functional space”(Langs et al., 2010). This kind of alignment may not produce meaningful results if the latent spaces themselves have a large disparity (Vos de Wael et al., 2020). On the other hand, generative models such as variational autoencoders (Kingma & Welling, 2013) can generate new data points and provide a bidirectional mapping between the data space and the latent space. Recently, a beta-variational autoencoder has been introduced for the automated discovery of interpretable factorized latent representations in images (Higgins et al., 2017), and has been used for disentangling resting-state fMRI activity in both adults (Kim et al., 2021) and fetus/neonates (Kim et al., 2023). Here, we introduce the use of a variational autoencoder to disentangle interpretable factors driving the variation in FC profile across both locations and subjects in a low-dimensional latent space. One toy example contrasting the comparison of FC profiles in vertex space and a low-dimensional latent space is provided in Figure 1A.

**Figure 1.**
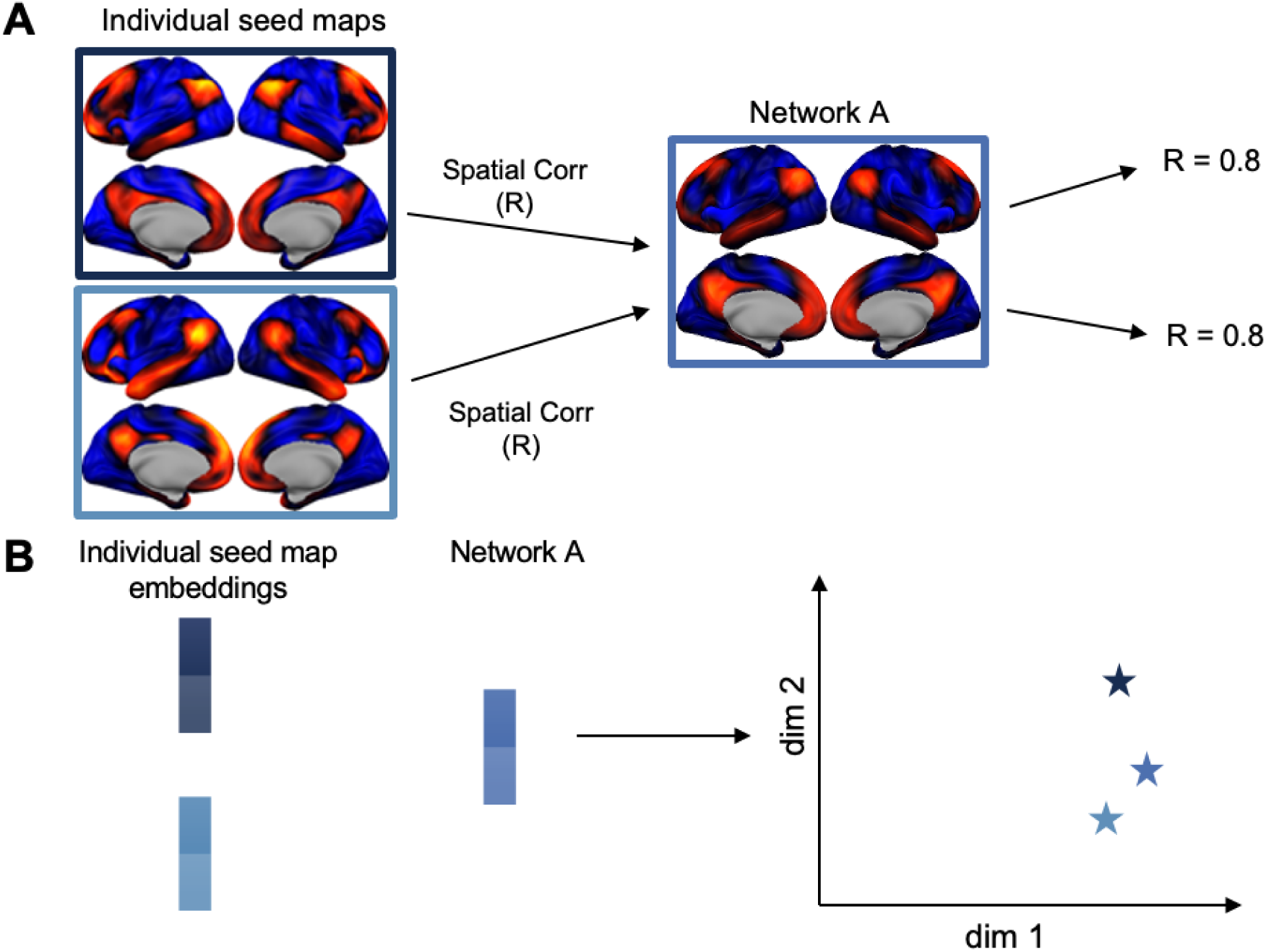
Comparing FC profiles in vertex space versus comparing FC profiles in the latent space. A) Vertex space: FC profiles from individual seed locations can have a similar spatial correlation to the FC profile of a network template even when they have subtle differences to each other. B) Latent space: the similarity between multiple FC profiles is approximated by the Euclidean distance between them in the latent space.

## 2 Methods

### 2.1 Neuroimaging Datasets

We used resting-state fMRI data from the Washington University 120 (WU120) (Power et al., 2014) for training the model weights to project the FC profiles from the original vertex space to a low-dimensional latent space. The WU120 dataset contains one resting-state session per subject for 120 subjects. Out of the 120 subjects from WU120, 100 subjects were selected as the training data, 10 as the validation data, and 10 as the test data. For additional analyses based on the pre-trained weights, we utilized the resting-state fMRI data from the Human Connectome Project (HCP, Rest1 and Rest2 scan sessions in 94 unrelated individuals) (Van Essen, Ugurbil, et al., 2012), the Midnight Scan Club (MSC, 10 scan sessions in 10 individuals)(Gordon, Laumann, Gilmore, et al., 2017) and Baby Connectome Project (BCP, 301 scan sessions in 178 individuals) (Howell et al., 2019) datasets. All datasets used are publicly available and the paths to data are provided in the “Data and Code Availability” section.

The adult datasets (all except BCP) were collected from young adult subjects (19-35 years) while they were asked to fixate on a center cross on the screen in a 3-Tesla MRI scanner. The baby dataset (BCP) was collected from infants to preschool children (8-60 months) during natural sleep. Procedures were then applied to normalize intensity, correct for motion in the scanner, and transform the data onto a standard 32k-fsLR surface (Van Essen, Glasser, et al., 2012). Furthermore, motion and other non-neuronal sources of artifact were mitigated with nuisance regression (including global signal regression) and bandpass filtering (Power et al., 2014) in the volume data before transforming to the surface space (adult datasets) and after transforming to the surface space (baby dataset). Further details on the acquisition and processing of the neuroimaging datasets and subject demographics are available in the Supplementary Materials.

### 2.2 Functional Connectivity Profiles

The FC profiles from each seed vertex were calculated as the Pearson’s correlation between the BOLD time series from that vertex to all cortical vertices in the left and right hemispheres (N = 59412 in the standard 32k-fsLR surface). FC profiles from a randomly sampled 10% of the vertices in each of the 100 subjects were used as the samples (N = 5942 × 100 = 594200) for training the model weights to map data from the vertex space (59412 dimensions) to the latent space to balance between the variability in the training samples and computational demand.

In addition, we also calculated the FC profiles from individual areas (N = 333 for the group-average adult parcellation (Gordon et al., 2016), N = 567-710 for Midnight Scan Club individual-specific parcellations (Gordon, Laumann, Gilmore, et al., 2017), and N = 326 for the group-average toddler parcellation), where each area consists of tens to hundreds of vertices. This is calculated with Pearson’s correlation between the average BOLD time series from each area and the BOLD time series from each of the 59412 vertices. FC profiles from functional networks (each with thousands of vertices) and the functional network prior (average FC profiles for functional networks across subjects) can be calculated with the same logic. Unless stated otherwise, the area parcellations and the FC profiles were visualized on a group-average brain surface in the standard 32k-fsLR space based on the MNI or Conte69 templates (Brett et al., 2002; Glasser & Van Essen, 2011).

### 2.3 The Variational Autoencoder Model

Autoencoders (AE) are neural networks designed to encode the input into a compressed representation, and then decode it back to a reconstructed input similar to the original one. The variational autoencoders (VAE) (Kingma & Welling, 2013) learn a distribution in the compressed representation. They are especially useful in obtaining a smooth, continuous latent space for generating new data, with the power to disentangle latent generative factors from images further enhanced with a higher weight on the Kullback-Leibler (KL)-divergence in the cost function with a hyperparameter β (Higgins et al., 2017). Unlike diffusion map embedding, AEs feature a encoder and decoder design for straightforward application to embed new data and reconstruct latent embeddings to the original vertex space. We adopted the same model architecture as described in prior research (Kim et al., 2021, 2023, 2024) with five convolutional layers and one fully-connected layer in the encoder, and one fully-connected layer and five convolutional layers in the decoder.

To take advantage of the convolutional layers in efficiently representing local patterns in images with few weights, we formatted the FC profiles in surfaces to 2D images. The geometric reformatting procedure was done in four steps. First, the FC profiles were mapped to the cortical surface using their coordinates in the 32k-fsLR mesh of the left and right hemispheres (32492 vertices per hemisphere with some of them empty due to the presence of the medial wall). Then, the surfaces in each hemisphere were inflated to a sphere using FreeSurfer (Fischl, 2012). After that, we used cart2sph.m in MATLAB to convert its Cartesian coordinates (x,y,z) to spherical coordinates (a,e), which reported the azimuth and elevation angles in a range from − π to + π and from − π/2 to + π/2, respectively. Lastly, we defined a 192×192 grid to resample the spherical surface with respect to azimuth and sin(elevation) such that the resampled locations were uniformly distributed at approximation (Figure 2A).

**Figure 2.**
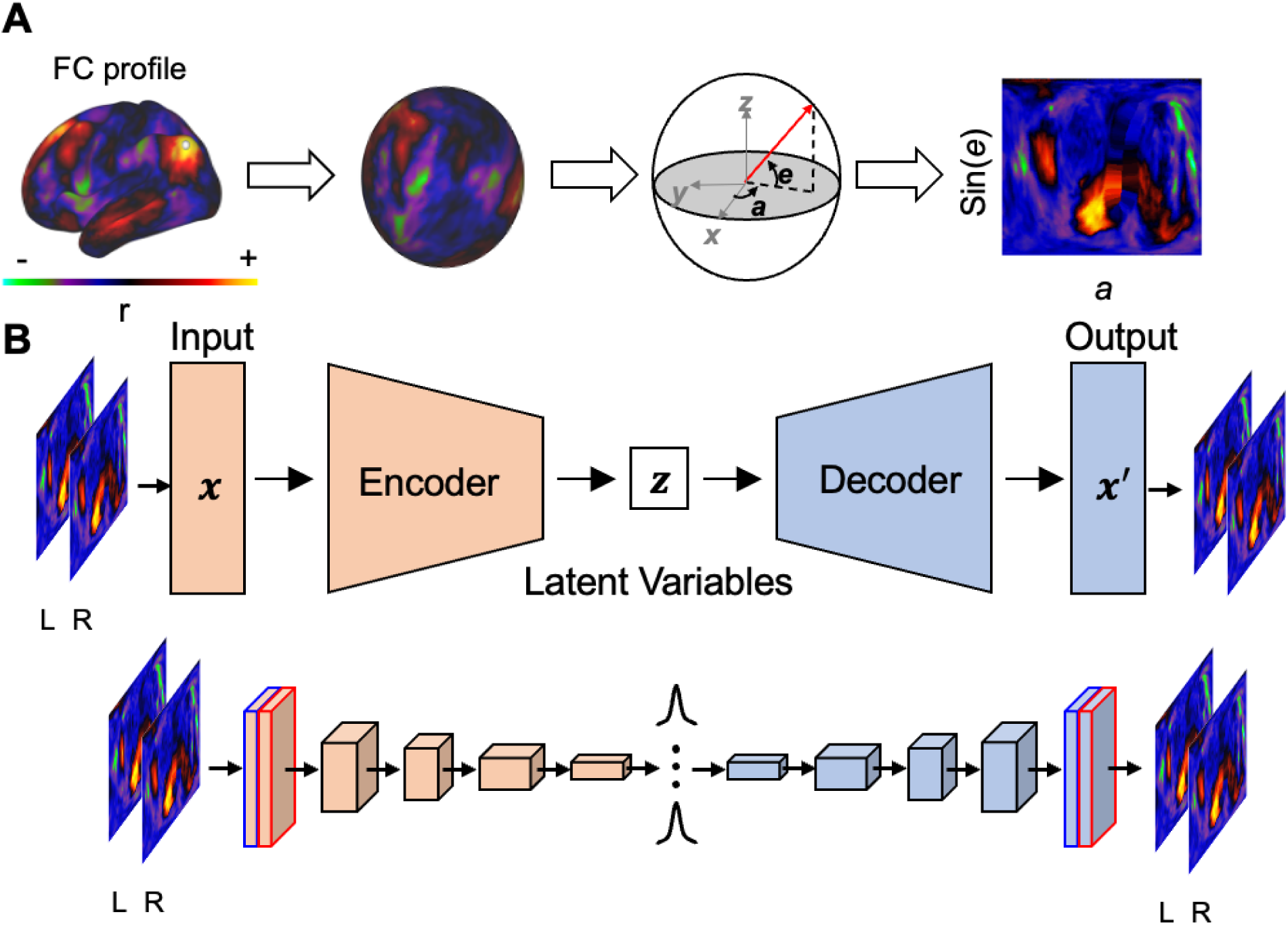
Geometric reformatting and the autoencoder model architecture. A) Geometric reformatting. The cortical distribution of fMRI activity is converted into a spherical surface and then to an image by evenly resampling the spherical surface with respect to sin(e) and a, where e and a indicate elevation and azimuth, respectively. B) Architecture of an autoencoder. An encoder network samples latent variables given an input image under the inference model while a decoder network generates a genuine input image from under the generative model. Both the encoder and decoder network contain 5 convolutional layers. Adapted from Kim, J., Zhang, Y., Han, K., Wen, Z., Choi, M., & Liu, Z. (2021). Representation learning of resting state fMRI with variational autoencoder. NeuroImage, 241, 118423. Copyright 2021 by Elsevier Inc.

The encoder transformed an FC profile (a pair of left and right hemisphere images formatted to two 192 x 192 grids) into a probabilistic distribution of N latent variables, where N is the number of latent dimensions. For visualization purposes, we use N = 2, although we have conducted additional experiments using different numbers of dimensions in Supplementary Materials (Section C). Each convolutional layer conducted linear convolutions followed by rectifying the outputs as described by (Nair & Hinton, 2010). The first layer utilized 8 × 8 convolutions on inputs from each hemisphere and combined the results. Subsequent layers, from the second to the fifth, applied 4 × 4 convolutions to this combined output. Circular padding was employed at the azimuth boundaries, while zero padding was used at the elevation boundaries. A fully connected layer applied linear weighting to generate the mean and standard deviation for the distribution of each latent variable. The decoder replicated this structure in reverse, reconnecting the layers to recreate the FC profile from a sample latent variable. The VAE model was optimized to reconstruct the input while constraining the distribution of every latent variable to be close to an independent and standard normal distribution. This is achieved through optimizing the encoding parameters, ϕ, and the decoding parameters, θ, to minimize the loss function below:

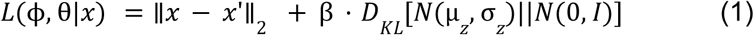

where *x* is the input data from both hemispheres, *x*’ is the reconstructed data, and *N*(µ_*z*_, σ_*z*_) is the posterior distribution, *N*(0, *I*) is the prior distribution. *D*_*KL*_ measures the KL divergence between the posterior and prior distributions, and β is a hyperparameter balancing the two terms in the loss function. A β < 1 places less emphasis on the KL divergence and focuses more on reconstruction, while a β > 1 places a higher emphasis on KL divergence, enforcing stricter regularization of the latent space.

Models were trained with stochastic gradient descent with a batch size of 128, initial learning rate of 1E-4, and 50 epochs with random data selection in each batch. An Adam optimizer (Kingma & Ba, 2014) was implemented, and the learning rate decayed by a factor of 10 every 20 epochs. Final hyperparameters including the number of latent dimensions and the beta value were determined by the trade-off between KL divergence and reconstruction loss on the validation data. The model was trained in Python 3.8 using PyTorch (v2.1.2+cu118) using a server with an NVIDIA A100 GPU (40 GB memory).

Additional details on the model design and hyperparameter tuning is provided in the Supplementary Materials (Section B). We also briefly explored alternative dimensionality reduction methods in the Supplementary Materials (Section D).

### 2.4 Quality of clustering by functional networks

One key feature of functional connectivity is that nodes within the same functional network tend to possess similar FC profiles (Yeo et al., 2011). The 286 out of 333 parcels in the Gordon parcellation (Gordon et al., 2016) were grouped into 12 functional networks. The additional (named “None” in the original network assignment) of 47 parcels in the low-SNR regions that cannot be confidently grouped into any of the functional networks were excluded from further analyses. We evaluated the segregation of FC profiles from different functional networks with the silhouette index (SI) (Rousseeuw, 1987; Yeo et al., 2011) on the group-average FC profiles or group-average latent embeddings. The SI is calculated as:

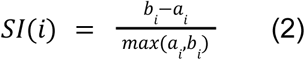

where *a*_*i*_ is the mean within-network distance, and *b*_*i*_ is the smallest mean between-network distance to alternative networks. A correlational distance measure was used for the FC profiles in the vertex space, whereas a Euclidean distance measure was used for the FC profiles in the latent space. A 95% confidence interval (95% CI) was calculated by bootstrapping the subject samples 1000 times.

## 3 Results

### 3.1 A Traversal Through the Latent Space Gives Rise to Systematic Variations in Reconstructed Functional Connectivity Profiles

To understand how the changes in the magnitude of each latent dimension affect the reconstructed FC profile appearance, we evenly divided one of the two latent dimensions (z_1_ and z_2_) while keeping the other latent dimension fixed at zero, and then back-project those latent embeddings to the vertex space using the VAE decoder. We observed patterns reminiscent of sensorimotor networks and association networks (Margulies et al., 2016; Sydnor et al., 2021), as well as task-positive to task-negative networks (Buckner et al., 2008; M. D. Fox et al., 2005; Raichle, 2015) (Figure 3A). Furthermore, somatomotor hand versus mouth and visual versus somatomotor network separations were also observable, suggesting the disentanglement of not only the coarse separation mentioned above but also fine details within the sensorimotor networks. Additionally, we obtained realistical FC profiles commonly observed across different functional networks for combinations of z_1_ and z_2_.

**Figure 3.**
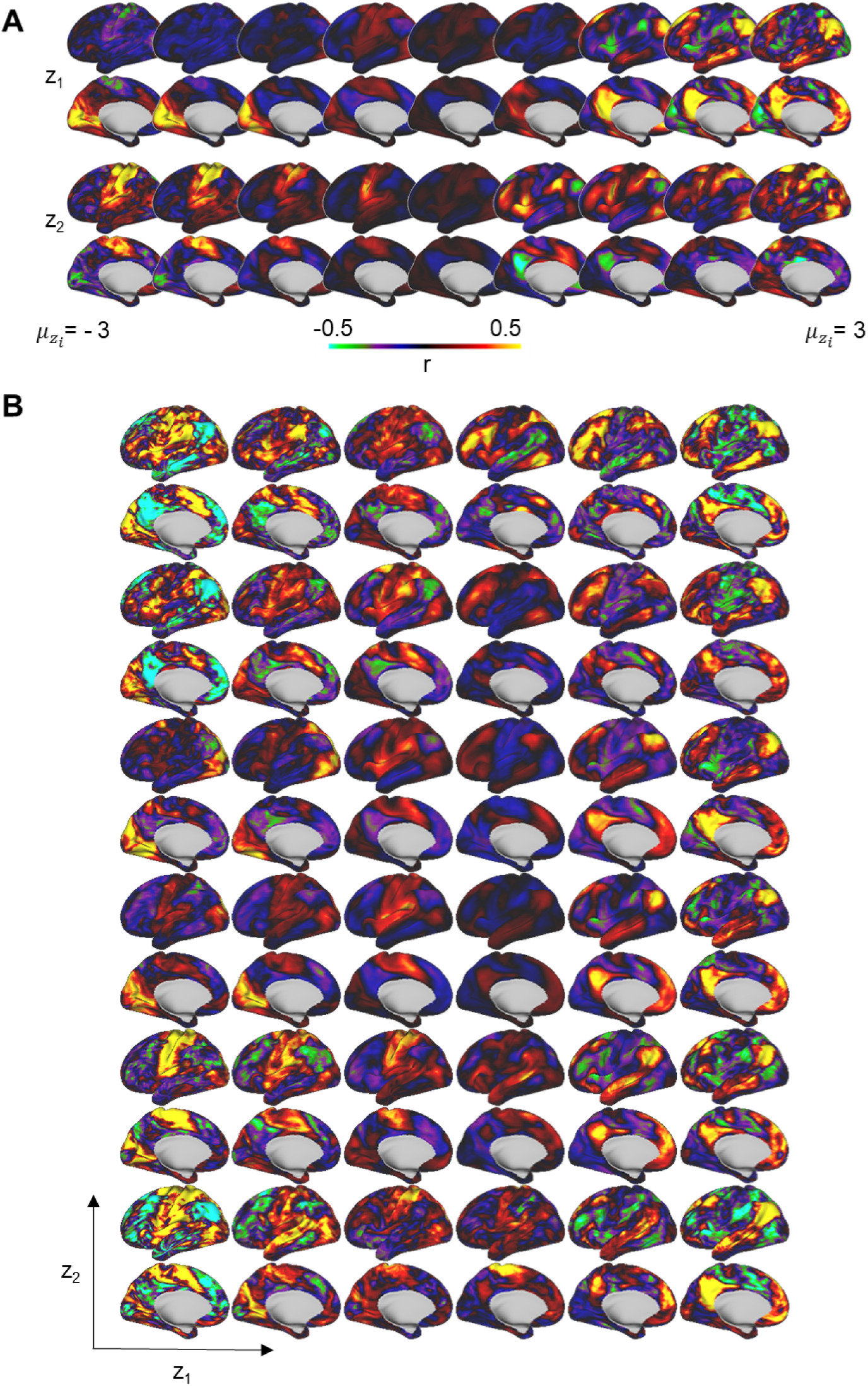
Reconstructed FC profiles obtained by traversing the latent space of the beta-VAE model. A) One latent dimension varies in equal steps from one end to the other, while the other dimension is fixed at zero. B) Grids representing different combinations of z_1_ and z_2_.

### 3.2 Functional Connectivity Profiles are Internally Coherent Within Functional Networks in Both the Vertex Space and the Latent Space

It hass been well established that FC profiles from the same functional networks tend to be similar (Buckner et al., 2013; Yeo et al., 2011). In this study, we validated this observation in the 94 unrelated subjects from the HCP dataset and evaluated whether similar patterns could be detected in the latent embeddings. When sorting the group-average FC profile from each of the 286 areas within the 12 networks defined by the Gordon parcellation (Gordon et al., 2016) according to the network order (Figure 4A), we observed that FC profiles within the same network appeared qualitatively similar (each row in Figure 4B is a flattened FC map from the 59412 cortical vertices). The relative similarity between the FC profiles (i.e., rows in Figure 4B) can be quantified using correlation distance (1-Pearson’s correlation). We found that within-network distances tend to be smaller than between-network distances (Figure 4C). On average, the FC profile of each area was more similar to those within the same network than to those in the closest alternative network (mean Silouette Index (SI) > 0) (Figure 4D). When projected into a two-dimensional latent space, the FC profiles form clusters that are closer together within the same functional networks, based on Euclidean distances (Figure 4E-F). On average, the FC embeddings from each area were closer to those within the same network than to those in the nearest alternative network (mean SI > 0). However, the mean SI was lower than previously observed in the vertex space, and some networks, such as the default mode network (red) were not distinctly separated from their closest alternative network. We will explore this observation further in the next section. Additionally, we found that the mean SI first increased and then decreased with the number of latent dimensions, potentially due to “curse of dimensionality” where all points are far apart at higher dimensions (Section C in the Supplementary Materials).

**Figure 4.**
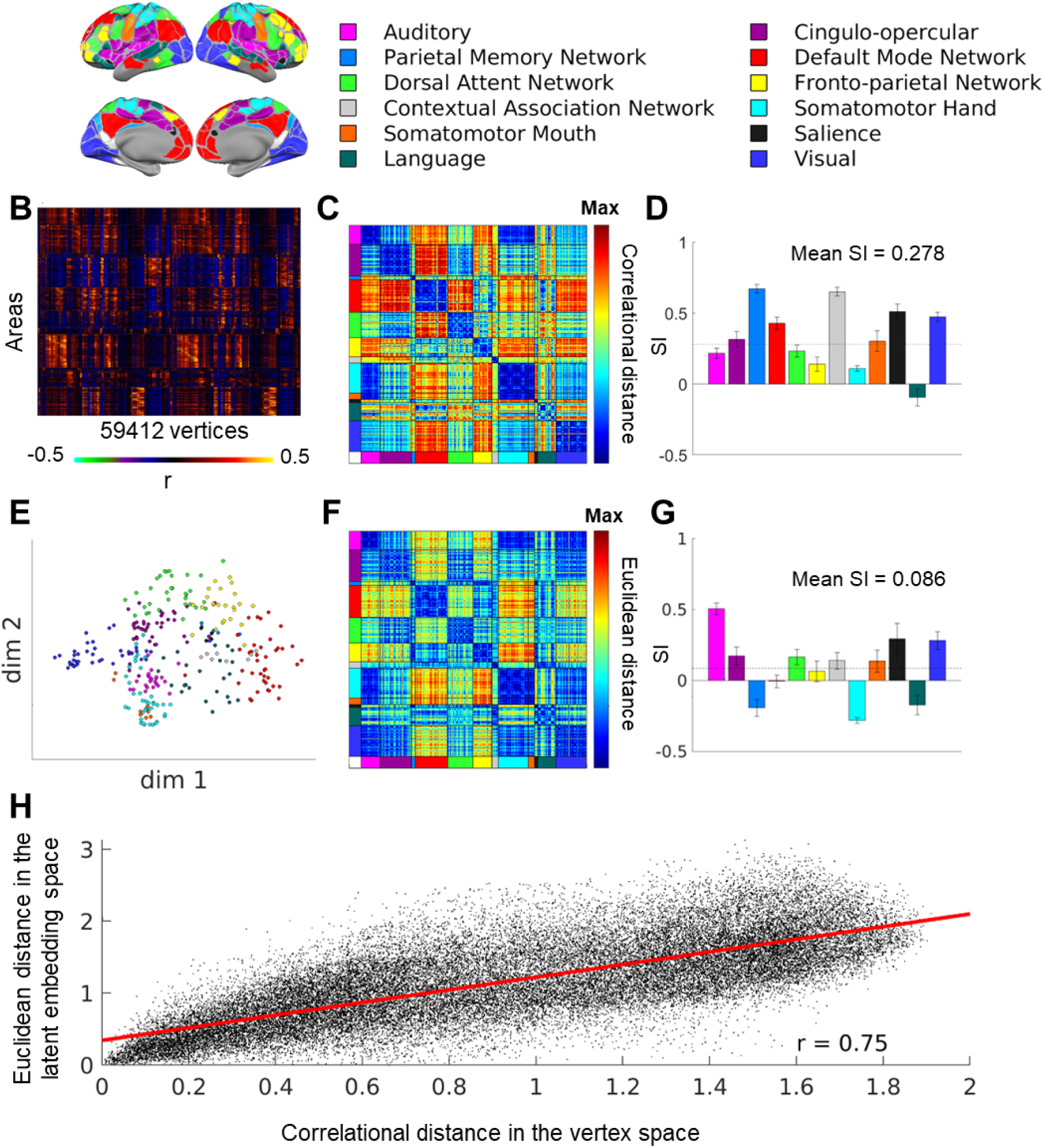
Separation of functional connectivity profiles by functional networks in the average of 94 HCP subjects. A) Gordon network assignments for 286 area parcels. B) The flattened FC profiles from each of the 286 area parcels. C) The mean correlational distance between each pair of the FC profiles in B. The dashed line shows the mean across all areas. D) The mean silhouette index for each functional network based on the correlation distance In C. E) The FC profile latent embeddings with two dimensions in VAE. Each circle represents each area parcel’s mean functional connectivity profile across Rest1 sessions of 94 subjects. F) The mean Euclidean distance between the latent embeddings of the average across 94 subjects. G) The mean silhouette index for each functional network based on the Euclidean distance in F. The dashed line shows the mean across all areas. H) A scatter plot of the Euclidean distance in the latent space (F) and correlational distance in the vertex space (C).

### 3.3 Functional Connectivity Profiles in the Default Mode Network Were Separated into Sub-networks in the Low-dimensional Latent Space

In addition to the group average, we can also examine the variability across the functional networks across all area parcels from the 94 subjects in the HCP dataset (Figure 5). It appears that while FC profiles from some networks form relatively local and spherical clusters in the latent space (e.g. the retrosplenial temporal network (RTN), salience network (Sal) and parietal memory network (PMN)), other networks exhibit more complex shapes and are sometimes divided into multiple spherical clusters across all subjects (e.g. default mode network (DMN)). To further investigate whether these clusters correspond to biologically meaningful divisions, we applied k-means clustering to the FC profiles from areas belonging to the default mode network from all subjects, choosing k = 2 based on visual inspection (Figure 6A-B). The resultant cluster consists of data from similar area parcels across the subjects (Figure 6C), albeit with some variability (Figure 6D). Cluster 1 primarily includes parcels in the medial prefrontal cortex, the temporal cortex and the dorsolateral prefrontal cortex, while Cluster 2 is mainly composed of parcels in the inferior parietal cortex and the posterior cingulate cortex (Figure 6E). Subtle differences can be observed in the reconstructed FC profiles from the centroids of the two clusters (Figure 6F). The reconstructed FC profile from the centroid of Cluster 1 resembles the ventromedial, pregenual and parietal components of the default network, while the distribution of Cluster 2 mirrors the dorsolateral and retrosplenial components of the default network. Similar division of the default mode subnetworks has been identified in other studies (Akiki & Abdallah, 2019; Andrews-Hanna et al., 2010; Gordon et al., 2020; Lynch et al., 2024; Uddin et al., 2009).

**Figure 5.**
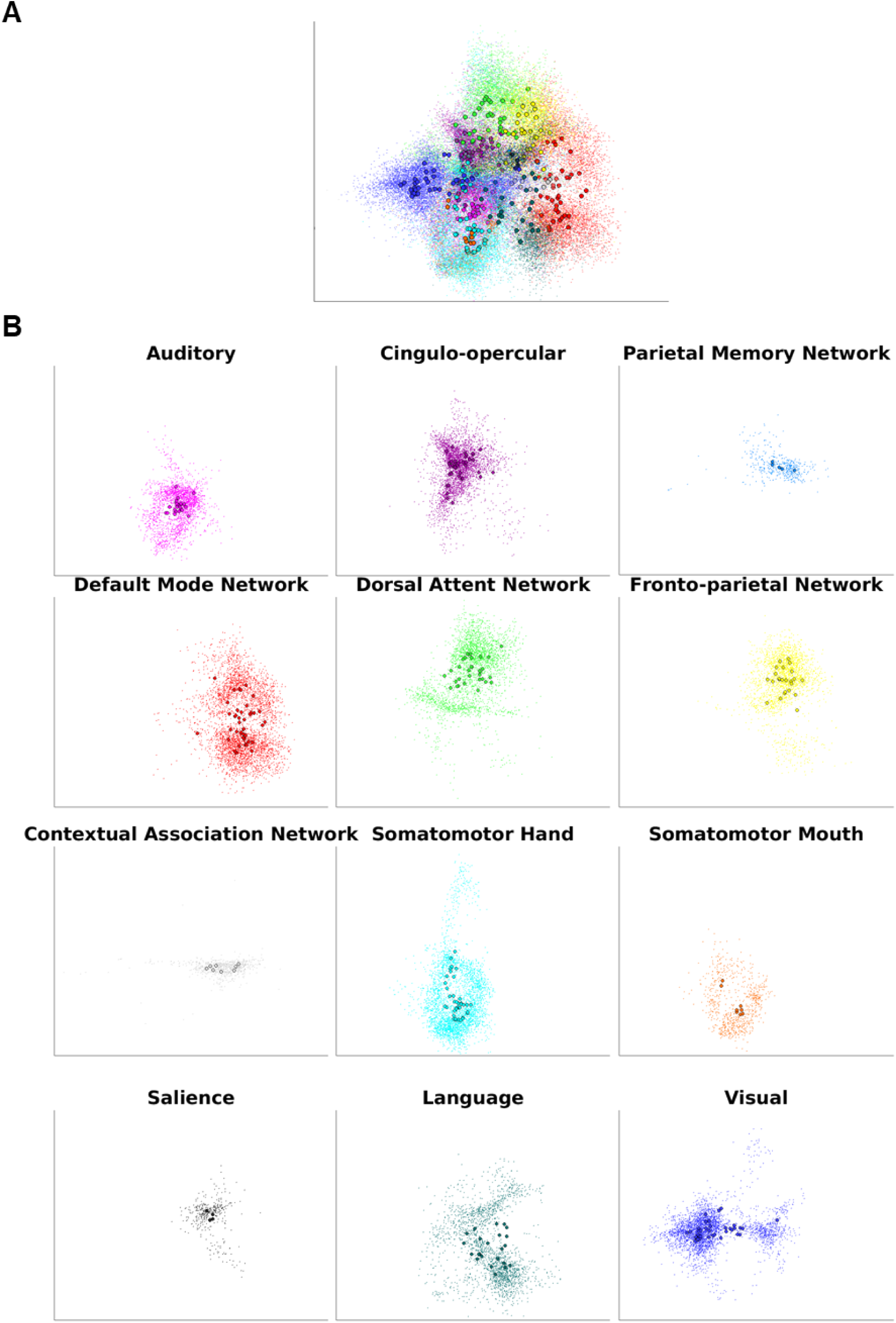
Functional connectivity profile embeddings from the Rest1 session for areas within the 12 Gordon networks across all 94 HCP subjects. A) All 12 networks. B) Each individual network.

**Figure 6.**
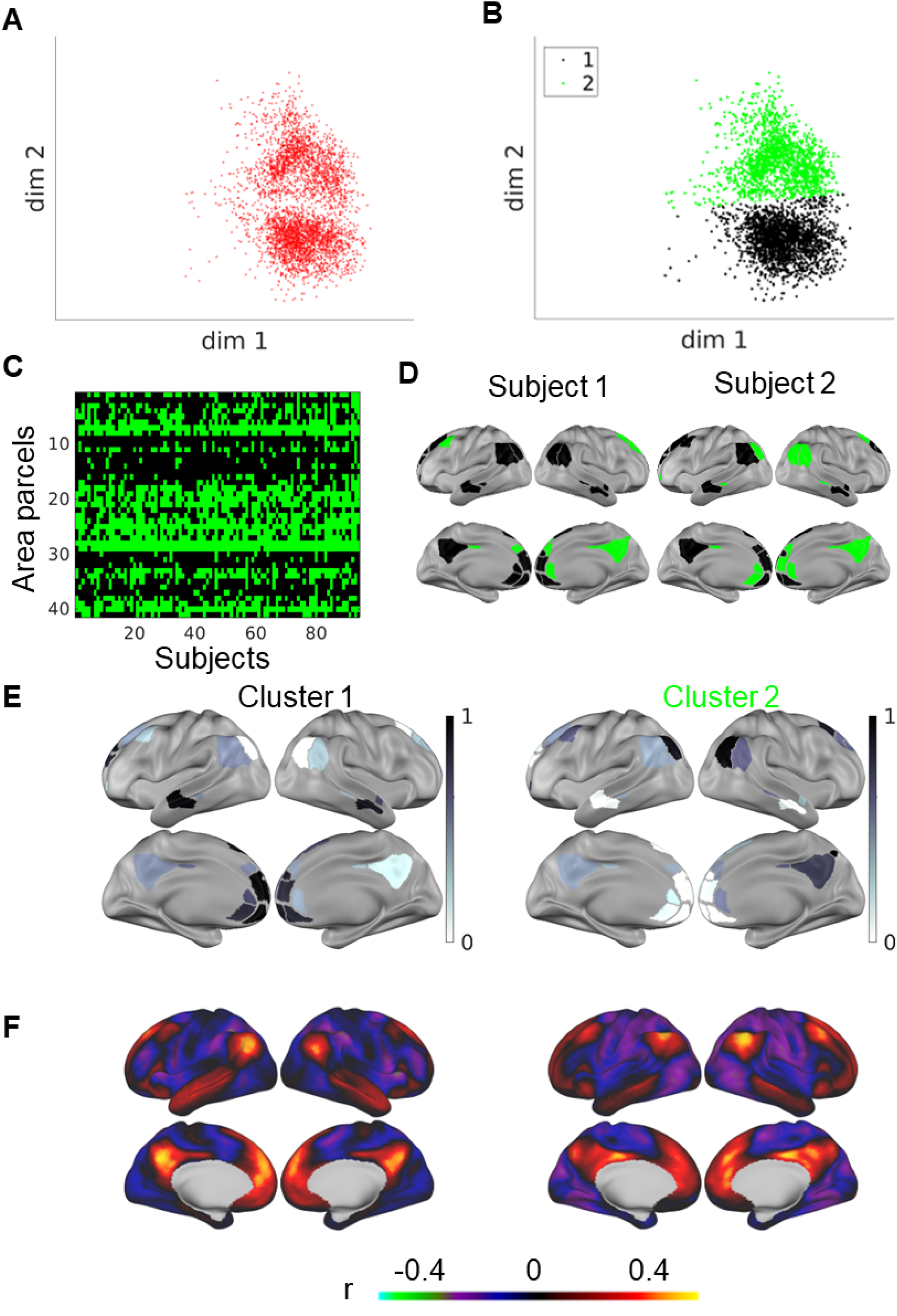
Sub-clusters within the DMN network in the latent space. A) DMN parcels across all subjects. B) Clustering the data in A into two sub-clusters with the k-means algorithm. C) The sub-cluster membership across parcels and subjects. D) The sub-cluster membership in two example subjects. E) The relative frequency of cluster membership across subjects (darker = more subjects) for the two sub-clusters. F) The reconstructed FC profiles from the centroids of the two sub-clusters. The axes scales were the same as Figure 5.

### 3.4 Interindividual Variability in Functional Connectivity Profiles From the Same Anatomically-matched Location was Evident in the Low-Dimensional Latent Space

Despite the largely consistent topography of functional networks across adult individuals (Damoiseaux et al., 2006; Gratton et al., 2018), interindividual differences in functional network assignment have been well documented in prior studies (Bijsterbosch et al., 2018; Braga & Buckner, 2017; Cui et al., 2020; Dworetsky et al., 2021, 2024; Glasser et al., 2016; Gordon, Laumann, Adeyemo, & Petersen, 2017; Gordon, Laumann, Adeyemo, Gilmore, et al., 2017; Gordon, Laumann, Gilmore, et al., 2017; Gratton et al., 2018; Kong et al., 2019; Kraus et al., 2021; Langs et al., 2016; H. Li et al., 2017; Seitzman, Gratton, et al., 2019; D. Wang et al., 2015). Here, we examine the position of the FC profile embedding from one example area parcel assigned to the “Somatomotor Hand” network based on the Gordon parcellation using group-average adult data (Gordon et al., 2016). The variability of FC profiles from this example parcel was substantial (Figure 7, the scales of the axes were the same as Figure 5-6). Additionally, the relative positions of the embeddings for different subjects were qualitatively consistent across two resting scan sessions (Rest1 and Rest2) in the VAE latent space (Figure 7). Visualization of the original FC profiles before projection to this low-dimensional latent space revealed that the two subjects at the extremes of the distribution demonstrated very distinct FC profiles: one resembling the “Dorsal Attention” network and the other “Somatomotor Hand” network (Figure 7).

**Figure 7.**
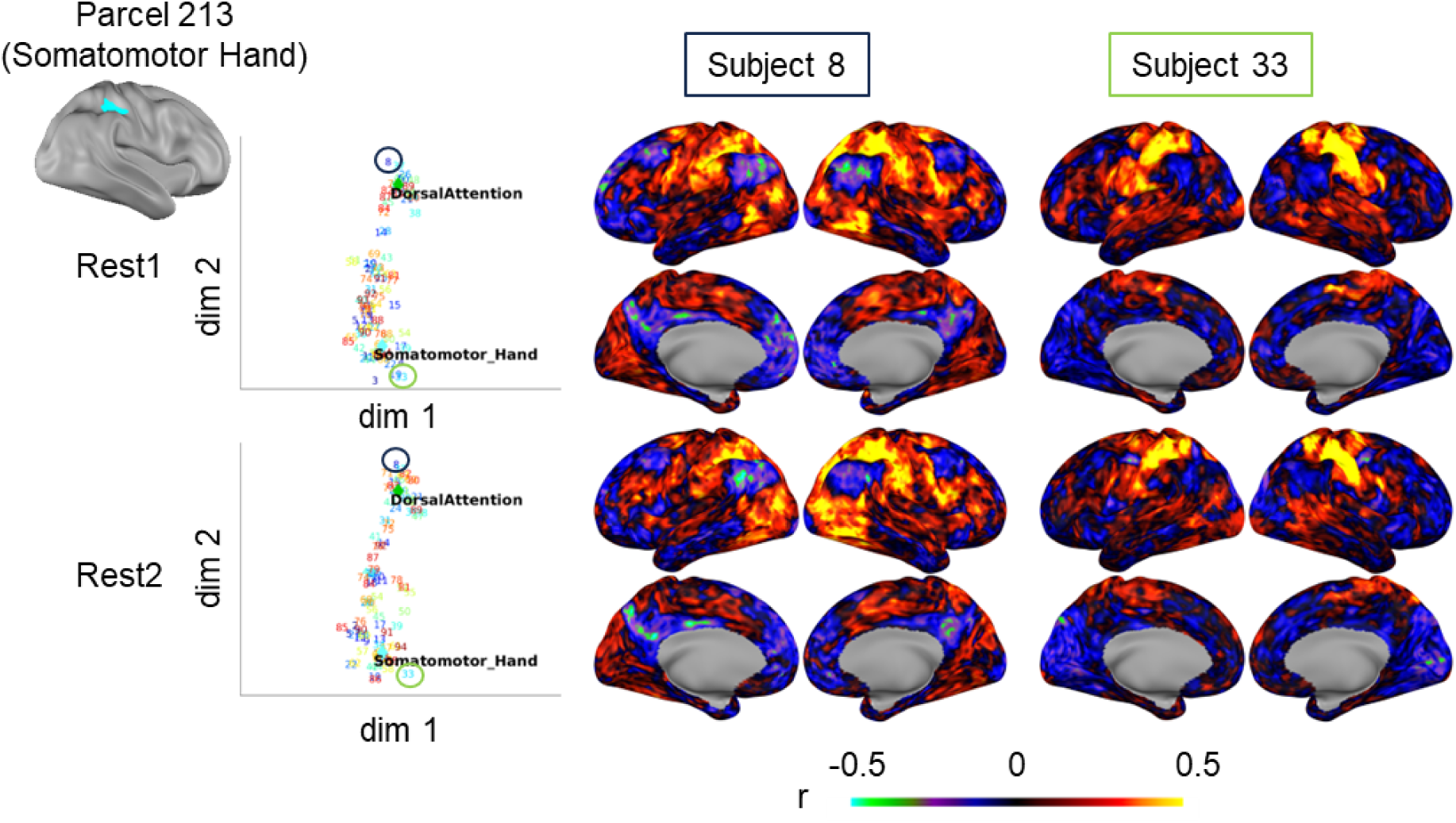
Interindividual variability of an example parcel across sessions across 94 HCP subjects in the VAE latent representation. Each data point is a subject indicated by the numbers 1 to 94. The FC profiles from parcel 213 for subject 8 and subject 33 in Rest1 and Rest2 were shown as an example. The “Somatomotor_Hand” and “DorsalAttention” markers were from Lynch et al. 2024. The axes scales were the same as Figure 5.

### 3.5 Low-dimensional Latent Representations of Functional Connectivity Profiles Enable Comparisons across Area Parcels, Sessions, Subjects and Populations

It is common for neuroimaging analyses to be conducted at the level of areas for biological interpretability (Petersen et al., 2024). However, different atlases exist for definitioning cortical areas, varying from 100-1000 areas across both hemispheres (Arslan et al., 2018; Craddock et al., 2012; Gordon et al., 2016; Schaefer et al., 2018; Shen et al., 2013). In addition to the lack of consensus in area definition, recent work has demonstrated that the optimal definition of areas may vary across individuals (Glasser et al., 2016; Gordon, Laumann, Gilmore, et al., 2017; Kong et al., 2021) and across the lifespan (Han et al., 2018; Myers et al., 2024; Tu, Myers, et al., 2024). Despite the variation in size and anatomical location of these parcels, we can obtain their FC profiles based on their functional connectivity to each vertex in the cerebral cortex. Subsequently, mapping those FC profiles to the low-dimensional latent space follows the previously described procedures straightforwardly. We mapped the distribution of all areas across all scan sessions in all subjects in two young adult datasets (HCP and MSC) and one baby dataset (BCP). The specific area and network assignments were reproduced in Supplementary Materials (Section A.6).

We found that the general compartments occupied by the major network divisions seem to be consistent across adult and baby datasets (Figure 8A/C/E). However, the baby dataset (8-60 months) had a different density distribution of points (Figure 8E). Additionally, the network priors calculated from the average of a group of highly sampled adult individuals for stereotypical appearance of FC in each network (Supplementary Figure A.6) (Lynch et al., 2024) also fall within the margins of the expected network distributions from the datasets (Figure 8B/D/F).

**Figure 8.**
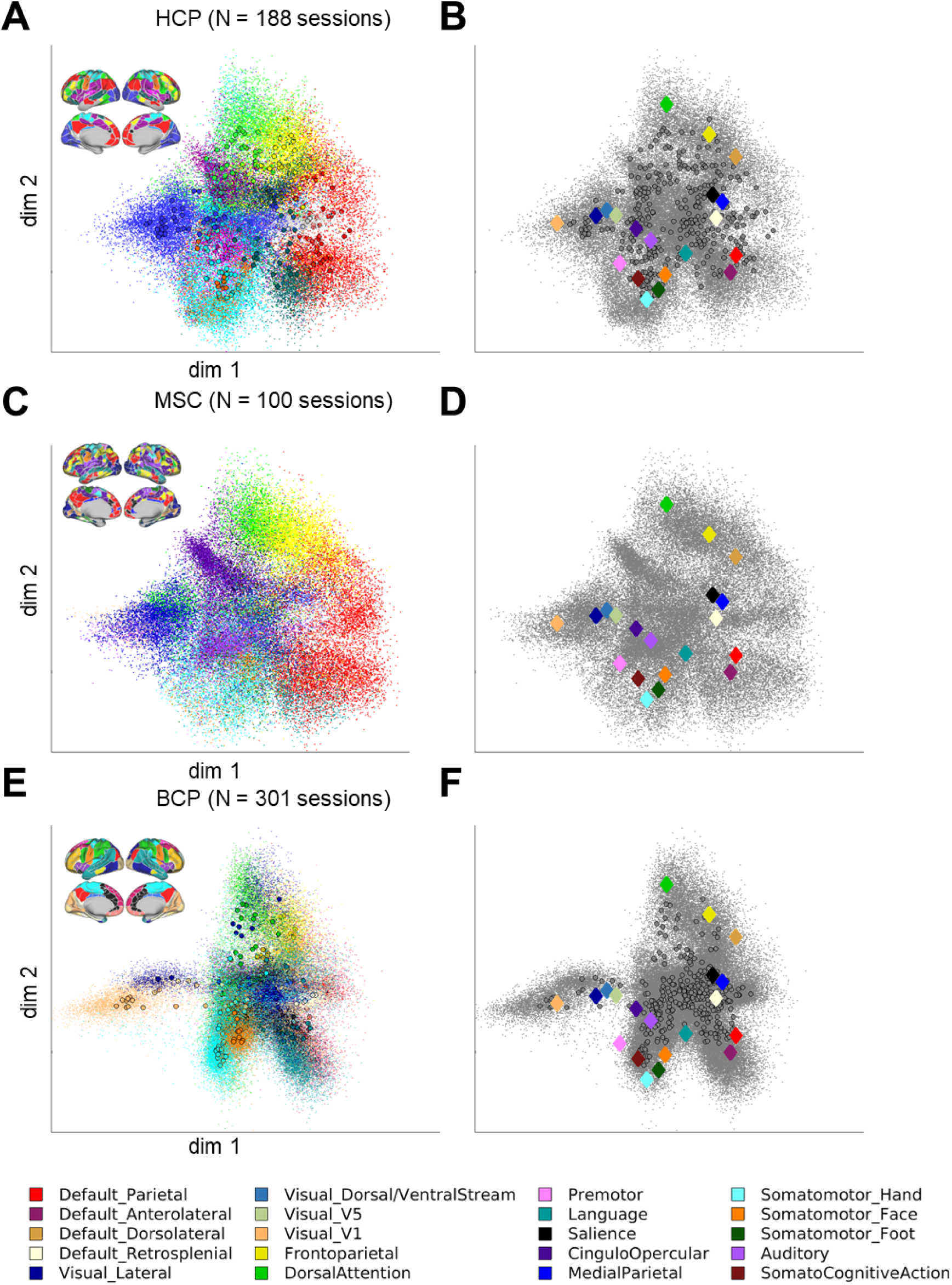
Distribution of functional connectivity profiles from different parcels in different cohorts. A) FC profiles from 286 area parcels (group-average adult parcellation) in 94 subjects in the HCP dataset with 2 scan sessions each. The network legend for the colors were the same as Figure 4A. C) FC profiles from 567-710 individual-level area parcels in 10 highly-sampled adult individuals with 10 scan sessions each. The area parcels displayed on the brain is from one example subject, with the area parcels and network legend for all subjects in the Supplementary Materials (Section A.6). E) 326 area parcels (group-average toddler parcellation) in 301 mixed longitudinal and cross-sectional sessions from 178 babies aged at 8-60 months, with the network legend in the Supplementary Materials (Section A.6). B/D/F is the same as A/C/E but without the network colors and with the overlay of network priors from Lynch et al. 2024.

We further quantified this difference in the distribution of FC profile embeddings across datasets by calculating the probability density function of the distributions (Figure 9). When comparing the baby (BCP) distribution (Figure 9B) with the young adult (HCP) distribution (Figure 9A), the BCP distribution was much denser at the center (Figure 9D), potentially attributable to an overall weaker FC (Figure 3). On the other hand, the difference in the FC profile embedding distributions in the two young adult datasets (HCP and MSC) (Figure 9C & 9E) was much smaller.

**Figure 9.**
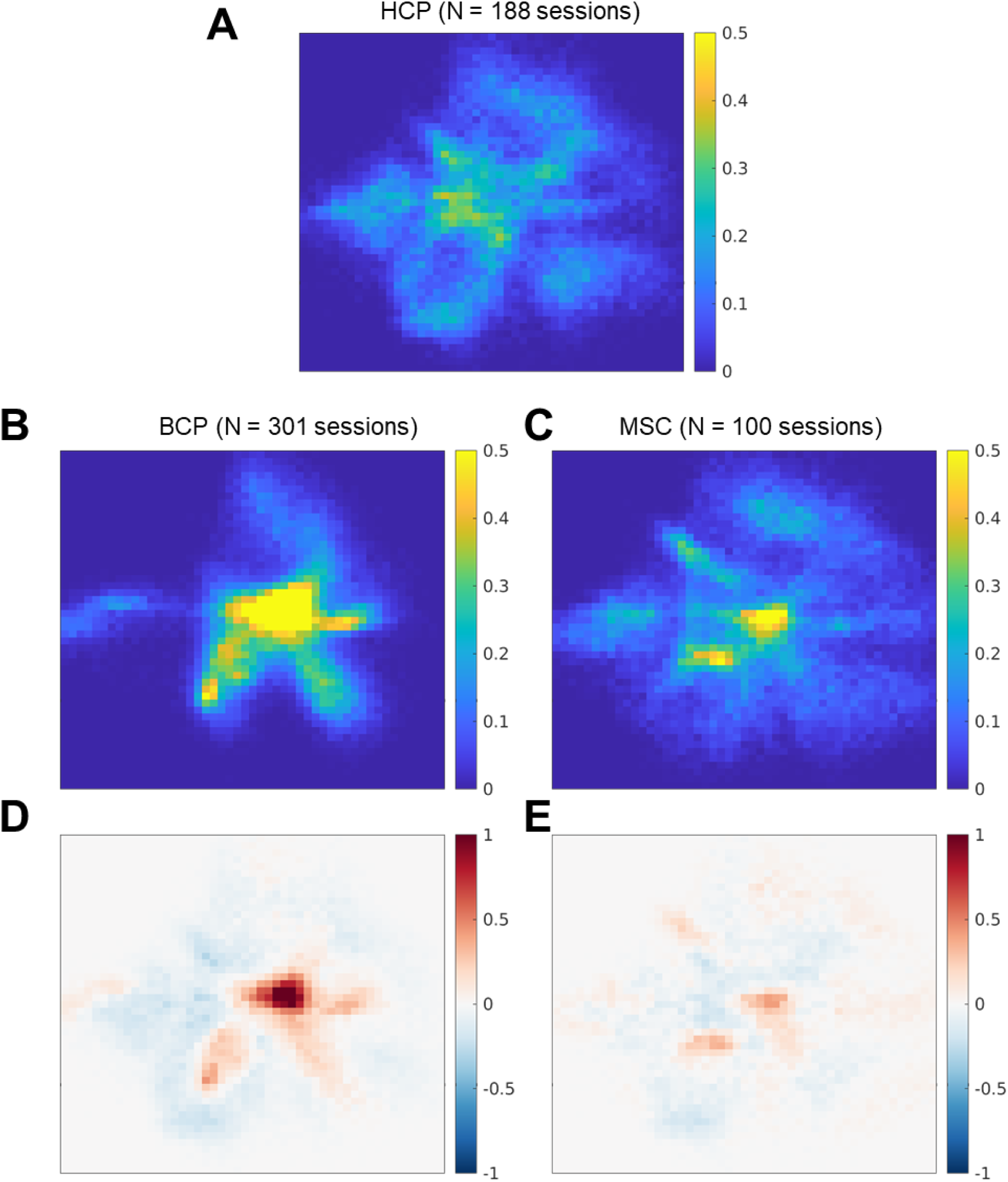
Probability density function in FC profile embedding distribution in the latent space across different cohorts. A) Probability density for the 188 sessions from the HCP data. B) Probability density for the 301 sessions from the BCP data. C) Probability density for the 100 sessions in the MSC data. D) Difference between B and A. E) Difference between C and A.

## 4 Discussion

### 4.1 A Low-dimensional Latent Space Facilitates Simultaneous Comparison of FC Profiles across Locations and Scans

Each point in the latent space represents a FC profile or a spatial connectivity map to all cortical vertices. The distribution of points characterizes the global functional connectivity pattern in each subject/group, which can vary between babies and adults (Figure 8-9). Neighboring points in the latent space exhibit coherent spatial patterns and form clusters reminiscent of functional networks (Figure 3-4). On the other hand, the anatomical positions on the brain are uncoupled from the positions in the latent space (Figure 6). Therefore, similar to other techniques that compare FC independent of the underlying anatomy (Guntupalli et al., 2018; Haxby et al., 2020; Langs et al., 2016), we group nodes (vertices, areas, etc.) together based on their similarity of FC profiles regardless of their anatomical locations. This approach is thus ideal for examining large-scale functional networks, which may originate from anatomically isolated compartments.

There are two advantages that our current method has over existing methods of FC embedding (Guntupalli et al., 2018; Haxby et al., 2020; Langs et al., 2016). First, the variational autoencoder is a generative model that offers a straightforward bidirectional mapping between the data space and the latent space, which allows for the visual assessment of example profiles from points in the latent space (Figure 6). Secondly, our approach facilitates easy projection of out-of-sample data with pre-computed weights, rather than first finding a data-driven embedding space for each subject and then aligning them. This simplifies the comparison across subjects and cohorts with minimal computational effort. To provide a more intuitive example, with a simple function call (see Code Availability), we were able to project the FC profiles from individual areas (approximately 600 per subject) in 100 sessions (10 MSC individual subjects with 10 sessions each) in 523 seconds on a Linux machine equipped with two AMD EPYC 7713 64-core Processors, providing 128 cores and 256 threads, with a base clock speed of 2.56 GHz.

### 4.2 Community Detection in a Shared Latent Space Naturally Establish Correspondence and Reduce Computational Demand

One key pursuit of system neuroscience is to group individual cortical areas into functional networks or communities across states, individuals, and the lifespan (Betzel et al., 2014; Dworetsky et al., 2021; Gordon, Laumann, Adeyemo, & Petersen, 2017; Grayson & Fair, 2017; Kong et al., 2019; Mitra et al., 2017; Muldoon & Bassett, 2016; Puxeddu et al., 2020; Sun et al., 2023; Tagliazucchi et al., 2013; D. Wang et al., 2015) based on their functional connectomes (i.e. area-to-area functional connectivity). A key hurdle for examining those communities is the establishment of correspondence across scan sessions. Prior work has attempted this by detecting communities in individuals and then matching their topographical overlap either through visual inspection (Gordon, Laumann, Gilmore, et al., 2017) or using a Hungarian matching algorithm (Kuhn, 1955) to relabel the networks such that the spatial agreement between corresponding networks is maximized (Langs et al., 2016; Yeo et al., 2014). Alternatively, a multilayer network can be constructed by linking multiple functional connectomes as layers (Bassett et al., 2011; Betzel et al., 2019; Puxeddu et al., 2020), where community detection methods are then applied to the entire multilayer network. However, neither approach can be applied to functional connectomes when the area definitions differ across individuals. At the same time, areas optimized for individuals (Gordon, Laumann, Gilmore, et al., 2017; Kong et al., 2021; Laumann et al., 2015; M. Li et al., 2019) or populations (Han et al., 2018; Myers et al., 2024; Scheinost et al., 2016; Shi et al., 2018; Tu, Myers, et al., 2024; F. Wang et al., 2023) have gained popularity over the years. Moreover, anatomically matched regions across individuals can still vary in their functional properties (Dworetsky et al., 2021; Gordon, Laumann, Adeyemo, & Petersen, 2017; Guntupalli et al., 2018; Haxby et al., 2020; Langs et al., 2016; Mueller et al., 2013).

Clustering in a low-dimensional latent space instead of the original data space can be a computationally efficient (Zhakubayev & Hamerly, 2022) way of solving the correspondence across individuals (Langs et al., 2016; Wen et al., 2024). The FC profiles from different individuals naturally acts as a prior for each other to compensate for the short acquisition time in each scan compared to the precision neuroimaging data (Allen et al., 2022; Gordon, Laumann, Gilmore, et al., 2017; Laumann et al., 2015; Lynch et al., 2024; Poldrack et al., 2015).

### 4.3 Challenges and future directions

Our training data represents a small sample with a narrow demographic profile. The training dataset could be expanded to incorporate diverse demographics, acquisition parameters, developmental stages, etc. Additionally, our current study focused on resting-state functional connectivity with a passive viewing or sleep paradigm, but recent studies suggest that a more naturalistic viewing paradigm might better emphasize inter-individual variability (Vanderwal et al., 2017). Future studies could incorporate FC collected in those more naturalistic settings.

To develop a general-purpose model for efficient bidirectional transformation between the spatial FC profiles in the latent space and the vertex space, we chose to directly embed the spatial FC profiles instead of the spatiotemporal maps (Kim et al., 2021, 2023, 2024), even though this overlooks the rich temporal information within fMRI timeseries. Additionally, unlike other factor analysis methods that explore harmonic modes/eigenmodes in the brain based on geometric anatomy (Atasoy et al., 2016, 2018; Pang et al., 2023), the VAE latent space lacks interpretability. Therefore, our model serves as a descriptive/phenomenological model to facilitate exploratory analyses in scientific discovery for functional connectivity analyses only, rather than a mechanistic model to understand the structural basis of functional connectivity.

## 5 Conclusion

The current study demonstrates the application of a variational autoencoder (VAE) to transform functional connectivity profiles from the original vertex space to a low-dimensional latent space. Unlike traditional non-linear dimensionality reduction methods like diffusion map embedding (Langs et al., 2010; Margulies et al., 2016), Laplacian Eigenmap (Haak et al., 2018; Pospelov et al., 2021), isomap (Pospelov et al., 2021) and local linear embedding (Pospelov et al., 2021), the variational autoencoder naturally offers a bidirectional mapping to and from the low-dimensional latent space. This bidirectionality allows the VAE to generate systematically varying FC profiles from samples in the latent space, providing valuable biological intuition. Additionally, by utilizing pre-trained weights from an independent young adult dataset (WU120), the VAE can generate meaningful latent representation without the need for individual-specific latent space computation and post-hoc alignment (Haxby et al., 2020; Langs et al., 2016; Nenning et al., 2020; Vos de Wael et al., 2020). This latent representation successfully capture the organization of FC profiles into functional networks and highlights the interindividual variability in FC profiles from matching anatomical locations. Our approach offers a robust dimensionality reduction technique for FC profiles across various area parcels, sessions, subjects and populations, including abstract summary measures such as network priors based on population averages (Lynch et al., 2024). This work could enhance visualization and exploratory analysis for precision functional mapping, providing insights into both individual and group differences in functional connectivity organization across the brain.

## Data and Code Availability (mandatory unless there is no data or code used)

Dataset WU120 is available at: https://openneuro.org/datasets/ds000243/versions/00001/file-display/00001

Dataset HCP is available at: https://www.humanconnectome.org/study/hcp-young-adult/document/1200-subjects-data-release

Dataset MSC is available at: https://openneuro.org/datasets/ds000224/versions/00001

Dataset BCP is available at: https://nda.nih.gov/edit_collection.html?id=2848

The Pytorch implementation for the beta-VAE can be found at https://github.com/cindyhfls/Tu-2025-VAE_FC_embedding, which was adapted from: https://github.com/libilab/rsfMRI-VAE. Pre-trained model weights can be found at https://huggingface.co/cindyhfls/fcMRI-VAE.

Visualization is conducted through Connectome Workbench (https://www.humanconnectome.org/software/connectome-workbench) and custom MATLAB scripts (https://github.com/cindyhfls/MATLAB_BrainParcelVisualizationFunctions)

## Author Contributions (mandatory)

JCT - Conceptualization, Methodology, Validation, Formal Analysis, Investigation, Data Curation, Writing - Original Draft, Writing - Review & Editing, Visualization and Project administration

JHK - Methodology, Investigation, Writing - Review & Editing, Project administration

PL - Methodology, Resources, Writing - Review & Editing.

BA - Resources, Data Curation, Writing - Review & Editing.

JSS - Methodology, Resources, Writing - Review & Editing.

JTE - Funding acquisition, Writing - Review & Editing.

ATE - Supervision, Writing - Review & Editing.

MDW - Supervision, Funding acquisition, Writing - Review & Editing.

## Funding (optional)

Funding for this work is provided through NIH research grants K99/R00 EB029343 and R01HD115540.

## Declaration of Competing Interests (mandatory)

The authors declare no competing interest relevant to this work.

## Acknowledgments

We would like to thank Dr. Evan M. Gordon for providing help with understanding the MSC dataset, Dr. Eric C. Leuthardt for providing computational resources for the current work and all participants and research personnel who helped with the collection, processing and curation of neuroimaging data.

## Supplementary Materials

### Section A. Neuroimaging Datasets

#### A.1 Washington University 120 (WU120) Data Acquisition and Processing Details

This dataset has been extensively detailed in prior descriptions (Power et al., 2014). In summary, data was obtained from 120 healthy young adults, with relaxed eyes-open fixation (60 females, average age = 25 years, age range = 19–32 years). Participants were right-handed, native English speakers recruited from the Washington University community. Screening via self-report questionnaire ensured no history of neurological or psychiatric diagnosis, nor head injuries resulting in more than 5 minutes of unconsciousness. All participants provided informed consent, and the study was approved by the Washington University School of Medicine Human Studies Committee and Institutional Review Board.

Structural and functional MRI data were obtained with a Siemens MAGNETOM Trio Tim 3.0-T Scanner and a Siemens 12-channel Head Matrix Coil. Structural imaging included a T1-weighted sagittal magnetization-prepared rapid acquisition gradient-echo (MP-RAGE) structural image was obtained [time echo (TE) = 3.08 ms, time repetition, TR (partition) = 2.4 s, time to inversion (TI) = 1000 ms, flip angle = 8°, 176 slices with 1 × 1 × 1 mm voxels]. Functional scan slices were aligned parallel to the anterior commissure–posterior commissure plane of the MP-RAGE and centered on the brain using an auto-align pulse sequence protocol available in Siemens software. This alignment corresponds to the Talairach atlas (Talairach & Tournoux, 1988).

For the fMRI data acquisition, subjects were instructed to relax while maintaining fixation on a black crosshair against a white background. Functional imaging used a BOLD contrast-sensitive gradient-echo echo-planar imaging (EPI) sequence (TE = 27 ms, flip angle = 90°, in-plane resolution = 4 × 4 mm). Full-brain EPI volumes (MR frames) of 32 contiguous, 4-mm-thick axial slices were obtained every 2.5 seconds. Additionally, a T2-weighted turbo spin-echo structural image (TE = 84 ms, TR = 6.8 s, 32 slices with 1 × 1 × 4 mm voxels) in the same anatomical planes as the BOLD images was also captured to augment atlas alignment. The fMRI acquisition used Anterior→Posterior (AP) phase encoding. The number of volumes collected per subject ranged from 184 to 724, with an average of 336 frames (14.0 min).

Functional images were first processed to reduce artifacts including (1) correction of odd versus even slice intensity differences due to interleaved acquisition without gaps, (2) head movement correction within and across runs, and (3) across-run intensity normalization to a whole-brain mode value of 1000 (Miezin et al., 2000). Each individual’s functional data was transformed into an atlas space using the MP-RAGE scan and resampled to an isotropic 3-mm atlas space (Talairach and Tournoux, 1988), using a single cubic spline interpolation (Lancaster et al., 1995).

Additional preprocessing mitigated high-motion frame effects in two iterations. The first iteration included: (1) demeaning and detrending, (2) multiple regression including whole-brain, ventricular cerebrospinal fluid (CSF), and white matter signals, and motion regressors derived by Volterra expansion and (3) a band-pass filter (0.009 Hz < f < 0.08 Hz). Temporal masks were created in this iteration to flag motion-contaminated frames, identified using framewise displacement (FD), calculated as the squared sum of the motion vectors (Power et al., 2012). Volumes exceeding FD > 0.2 mm and segments of fewer than 5 contiguous volumes were flagged for removal.

The data were then reprocessed in a second iteration, incorporating the temporal masks described above. This reprocessing was identical to the initial processing stream but ignored censored data. Data were interpolated across censored frames using least squares spectral estimation (Power et al., 2014) of the values at censored frames so that continuous data could be passed through the band-pass filter (0.009 Hz < f < 0.08 Hz) without contaminating frames near high motion frames. *Censored frames were ultimately ignored when calculating the functional connectivity profiles*.

Individual surfaces were generated from the structural images and functional data were sampled to surface space (Glasser et al., 2013). Following volumetric registration, left and right hemisphere anatomical surfaces were created from each subject’s MP-RAGE image using FreeSurfer’s recon-all processing pipeline (v5.0)(Fischl, 2012). This involved brain extraction, segmentation, white matter and pial surface generation, surface inflation to a sphere, and spherical registration of the subject’s “native” surface to the fsaverage surface. The fsaverage-registered surfaces were then aligned and resampled to a resolution of 164000 vertices using Caret tools (Van Essen et al., 2001) and down-sampled to a 32492 vertex surface (32k-fs_LR). Functional BOLD volumes were sampled to each subject’s individual “native” midthickness surface (generated as the average of the white and pial surfaces) using the ribbon-constrained sampling procedure available in Connectome Workbench (v0.84) and then deformed and resampled from the individual’s “native” surface to the 32k-fs_LR surface. The final time series were smoothed along the 32k-fs_LR surface using a Gaussian smoothing kernel (σ = 2.55 mm).

#### A.2 Human Connectome Project (HCP) Data Acquisition and Processing Details

Data for the Human Connectome Project (Young Adult) was collected at Washington University in St. Louis and the University of Minnesota. The participants, healthy adults aged between 22 to 35 years, underwent high-resolution T1-weighted (MP-RAGE, TR = 2.4s, voxel size = 0.7×0.7×0.7mm) and BOLD contrast-sensitive imaging (gradient echo EPI, multiband factor 8, TR = 0.72s, voxels = 2×2×2mm) using a custom Siemens SKYRA 3.0T MRI scanner equipped with a custom 32-channel Head Matrix Coil. Sequences with both left-to-right (LR) and right-to-left (RL) phase encoding were employed, with each participant completing a single run in each direction over two consecutive days, resulting in four runs in total, two for Rest 1 and another two for Rest 2 (Van Essen, Ugurbil, et al., 2012).

Functional data processing first followed the HCP minimally preprocessing pipeline (Glasser et al., 2013), which included the additional use of a field map for distortion correction (Cusack et al., 2003; Jezzard & Balaban, 1995) and otherwise similar steps as mentioned above. Then, additional preprocessing of the resting-state BOLD volume data was applied to remove non-neuronal sources of artifacts similar to the WU120 data, with the only differences in and an added low-pass filter at 0.1 Hz applied on the movement parameters before calculating FD to mitigate high-frequency respiration artifacts on FD estimates (Gratton, Dworetsky, et al., 2020). Frames with FD greater than 0.04 mm were flagged as “high-motion” based on the noise floor in FD traces. Instead of processing in two iterations as mentioned above for the WU120 data, the temporal mask excluding high-motion frames based on FD was used in the calculation of regression coefficients and linear interpolation before the application of bandpass filtering.

##### Censored frames were ultimately ignored when calculating the functional connectivity profiles

Following that, the preprocessed BOLD volumes were sampled to each subject’s individual “native” midthickness surface (generated as the average of the white and pial surfaces) using the ribbon-constrained sampling procedure available in Connectome Workbench and then deformed and resampled from the individual’s “native” surface to the 32k-fs_LR surface. The final BOLD time series was minimally smoothed with a Gaussian kernel (FWHM = 2mm, σ = 0.85).

#### A.3 Midnight Scan Club (MSC) Acquisition and Processing Details

The acquisition and processing of this dataset have been mentioned in elaborative details in a prior study (Gordon, Laumann, Gilmore, et al., 2017). Data were collected from ten healthy, right-handed, young adult subjects (5 females; age: 24-34) recruited from the Washington University community. The study was approved by the Washington University School of Medicine Human Studies Committee and Institutional Review Board.

Imaging for each subject was performed on a Siemens TRIO 3T MRI scanner over the course of 12 sessions conducted on separate days, each beginning at midnight. Structural MRI was conducted across two separate days. In total, four T1-weighted images (sagittal, 224 slices, 0.8 mm isotropic resolution, TE = 3.74 ms, TR = 2400 ms, TI = 1000 ms, flip angle = 8 degrees), four T2-weighted images (sagittal, 224 slices, 0.8 mm isotropic resolution, TE = 479 ms, TR = 3200 ms) were obtained for each subject. On ten subsequent days, each subject underwent 1.5 hr of functional MRI scanning beginning at midnight. In each session, we first collected thirty contiguous minutes of resting state fMRI data, in which subjects visually fixated on a white crosshair presented against a black background. Each subject was then scanned during the performance of three separate tasks: motor (2 runs per session, 7.8 min combined), incidental memory (3 runs per session, 13.1 min combined), and mixed design (2 runs per session, 14.2 min combined). Across all sessions, each subject was scanned for 300 total minutes during the resting state and approximately 350 total minutes during task performance. All functional imaging was performed using a gradient-echo EPI sequence (TR = 2.2 s, TE = 27 ms, flip angle = 90°, voxel size = 4 mm x 4 mm x 4 mm, 36 slices). In each session, one gradient echo field map sequence was acquired with the same prescription as the functional images.

Following that, the preprocessed BOLD volumes were sampled to each subject’s individual “native” midthickness surface (generated as the average of the white and pial surfaces) using the ribbon-constrained sampling procedure available in Connectome Workbench and then deformed and resampled from the individual’s “native” surface to the 32k-fs_LR surface. The final BOLD time series was smoothed with a Gaussian kernel (FWHM = 6mm, σ = 2.55).

#### A.4 Baby Connectome Project Acquisition and Processing Details

Details about the acquisition and processing of this dataset have been previously reported (Tu, Wang, et al., 2024). Full-term (gestational age of 37-42 weeks) infants free of any major pregnancy and delivery complications were recruited as part of the Baby Connectome Project (Howell et al., 2019). All procedures were approved by the University of North Carolina at Chapel Hill and the University of Minnesota Institutional Review Boards. Informed consent was obtained from the parents of all participants. In the final cohort used following fMRI data quality control, we retained 303 fMRI sessions from 178 individuals acquired during natural sleep. All MRI images were acquired on a Siemens 3T Prisma scanner with a 32-channel head coil at the University of Minnesota and the University of North Carolina at Chapel Hill during natural sleep without the use of sedating medications. T1-weighted (TR=2400 ms, TE=2.24 ms, 0.8 mm isotropic; flip angle = 8°), T2-weighted images (TR=3200 ms, TE=564 ms, 0.8 mm isotropic), spin echo field maps (SEFM) (TR=8000 ms, TE=66 ms, 2 mm isotropic, MB=1), and fMRI data (TR=800 ms, TE=37 ms, 2 mm isotropic, MB=8) were collected. A mixture of Anterior→Posterior (AP) and Posterior→Anterior (PA) phase encoding directions was used for fMRI acquisition in each session, but they were concatenated into one time series. A subset of data had a 720-ms TR.

Data processing used the DCAN-Labs infant-abcd-bids-pipeline (v0.0.22) which largely follows the HCP processing(Glasser et al., 2013) and the steps described previously for the ABCD dataset (Feczko et al., 2021). The processing steps were similar to the adult data except for the use of infant-specific MNI templates to better register the structural data. In addition, segmentation of the brain structures was conducted with Joint Label Fusion (JLF). The toddler-specific brain mask and segmentation were substituted to the Freesurfer (Fischl, 2012) pipeline to refine the white matter segmentation and guide the FreeSurfer surface delineation for each individual scan session of each subject.The native surface data were then deformed to the 32k fs_LR template via a spherical registration.

For functional data processing, a scout image (frame 16 in each run) was selected from the fMRI time series. The scout was distortion-corrected via spin-echo field maps, served as the reference for motion correction via rigid-body realignment (Feczko et al., 2021), and was registered to the native T1. Across-run intensity normalization to a whole-brain mode value of 10,000 was then performed. These steps were combined in a single resampling with the MNI template transformation from the previous step, such that all fMRI frames were registered to the infant MNI template. Manual inspection of image quality of structural and functional data was conducted to exclude sessions with bad data quality. fMRI BOLD volumes were sampled to native surfaces using a ribbon-constrained sampling procedure available in Connectome Workbench and then deformed and resampled from the individual’s “native” surface to the 32k-fs_LR surface. Additional steps to mitigate the non-neuronal sources of artifact including demean/detrend, nuisance regression and bandpass filtering was conducted on the surface data similar to the adults with a few minor differences: 1) gray matter signal based on the 91k grayordinates in the 32k-fs_LR surface substituted the whole-brain signal in the nuisance regression, 2) a respiratory notch filter (0.28-0.48 Hz) was applied to the motion parameter estimates before FD calculation to minimize the inflation of FD values by the perturbation of magnetic field from subject respiration in fast TR scans (Fair, 2020; Kaplan et al., 2022). The data were originally minimally spatially smoothed with a geodesic 2D Gaussian kernel (σ = 0.85 mm). A further smoothing with a geodesic 2D Gaussian kernel (σ = 2.40 mm) was applied to give a final effective smoothing of σ = 2.55 mm.

#### A.5 Parcellation Atlases Used to Define Areas and Functional Networks in Each Dataset

The area parcellation atlases used for MSC data (Supplementary Figure A.5A) and BCP data (Supplementary Figure A.5B) with their network assignments were reproduced below from the original publications.

**Supplementary Figure A.5.**
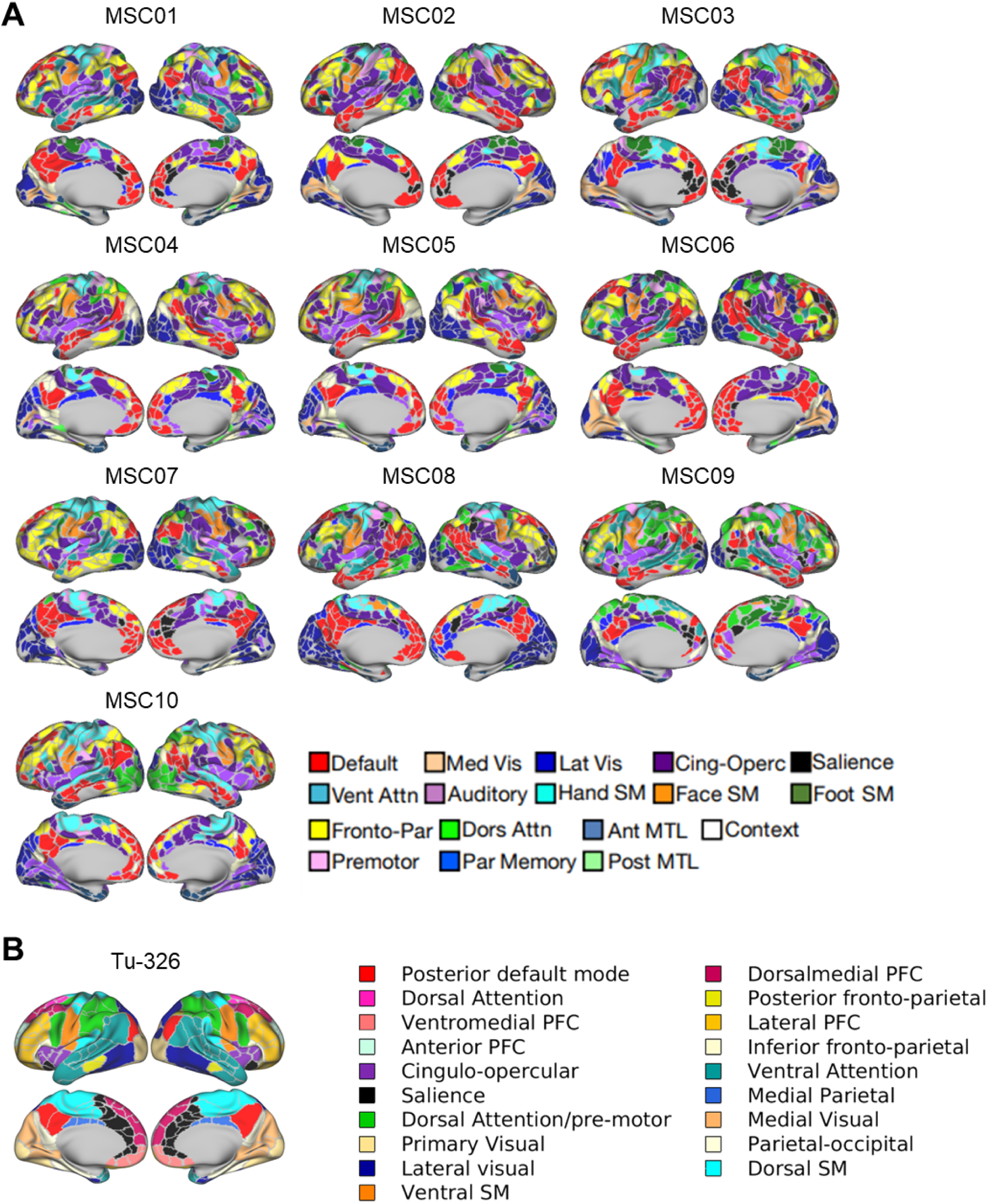
Parcellations used for defining areas and networks for the MSC and BCP datasets. A) Subject-specific parcellation and putative network identity of each parcel for each MSC subject. Reproduced from Gordon, E. M., Laumann, T. O., Gilmore, A. W., Newbold, D. J., Greene, D. J., Berg, J. J., … & Dosenbach, N. U. (2017). Precision functional mapping of individual human brains. Neuron, 95(4), 791-807. Copyright 2017 by Elsevier Inc. Default = Default mode. Med Vis = Medial Visual. Lat Vis = Lateral Visual. Vent Attn = Ventral Attention. SM = Somatomotor. Fronto-Par = Fronto-parietal. Dors Attn = Dorsal Attention. Ant MTL = anterior medial temporal lobe. Posterior MTL = posterior medial temporal lobe. Par Memory = Parietal memory. B) Group-average toddler parcellation.

#### A.6 Functional network priors from highly-sampled adult individuals

We reproduced the functional network priors figure calculated from 45 adult individuals (Lynch et al., 2024) from the original publication and indexed them to the locations on the 2D latent space.

**Supplementary Figure A.6.**
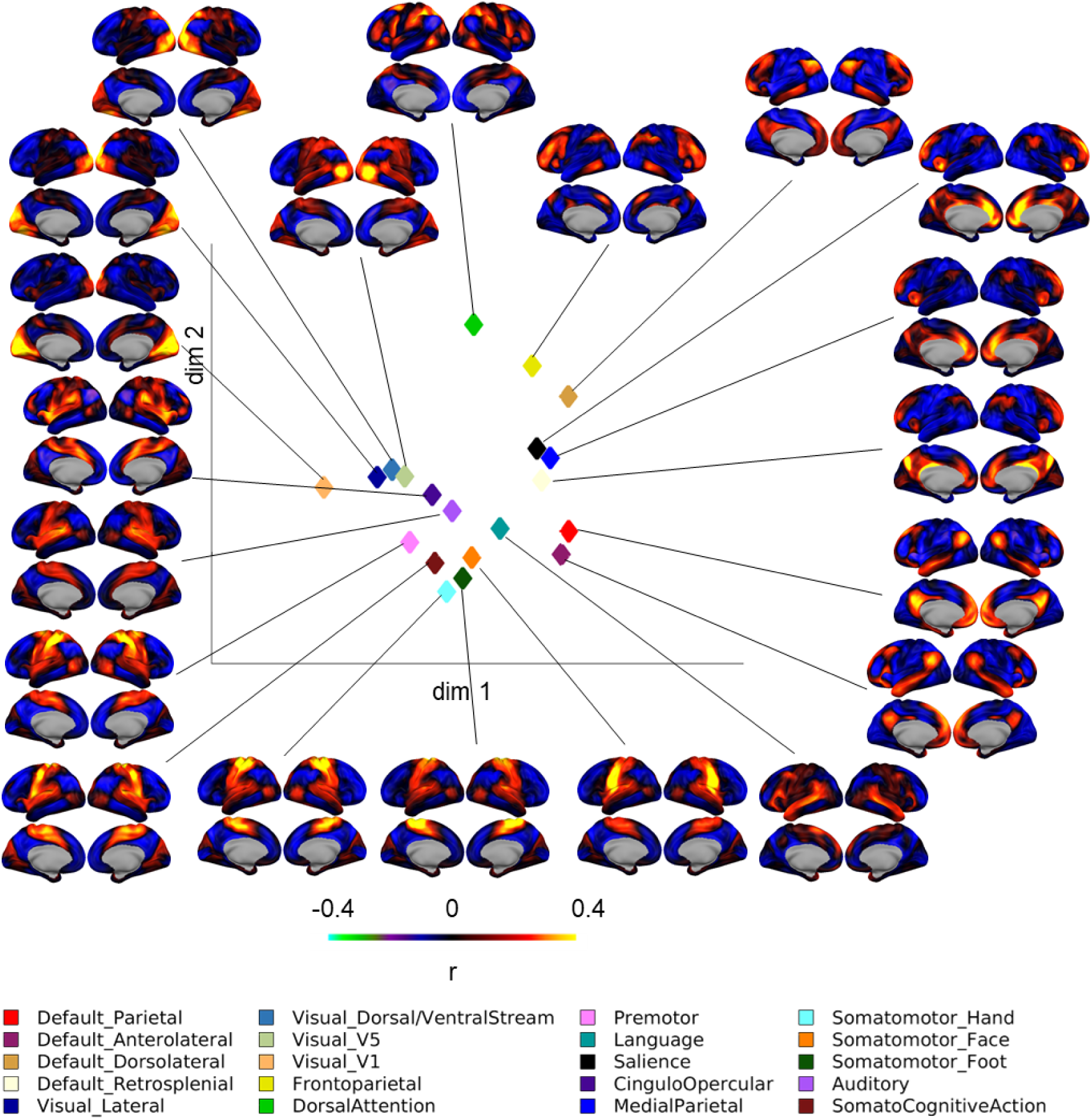
Functional priors and their location on the 2-D latent space. The functional network priors reproduced from data shared by Lynch, C.J., Elbau, I.G., Ng, T. et al. Frontostriatal salience network expansion in individuals in depression. Nature 633, 624–633 (2024).

### Section B. Model training details

#### B.1 Model architecture

The variational autoencoder consists of a total of 12 layers (Figure B.1). In the encoder network, the size of the output image of each layer (from left to right) is 96×96×64 (32 channels per hemisphere), 48×48×128, 24×24×128, 12×12×256, and 6×6×256; for the decoder network, 6×6×256, 12×12×256, 24×24×128, 48×48×128, and 96×96×64 (32 channels per image), from left to right. The dimension of latent variables is 256. The convolution operations are defined as 1: convolution (kernel size=8, stride=2, padding=3) with rectified nonlinearity, 2-5: convolution (kernel size=4, stride=2, padding=1) with rectified nonlinearity, 6: fully-connected layer with re-parametrization, 7: fully-connected layer with rectified nonlinearity, 8-11: transposed convolution (kernel size=4, stride=2, padding=1) with rectified nonlinearity, 12: transposed convolution (kernel size=8, stride=2, padding=3).

**Supplementary Figure B.1.**
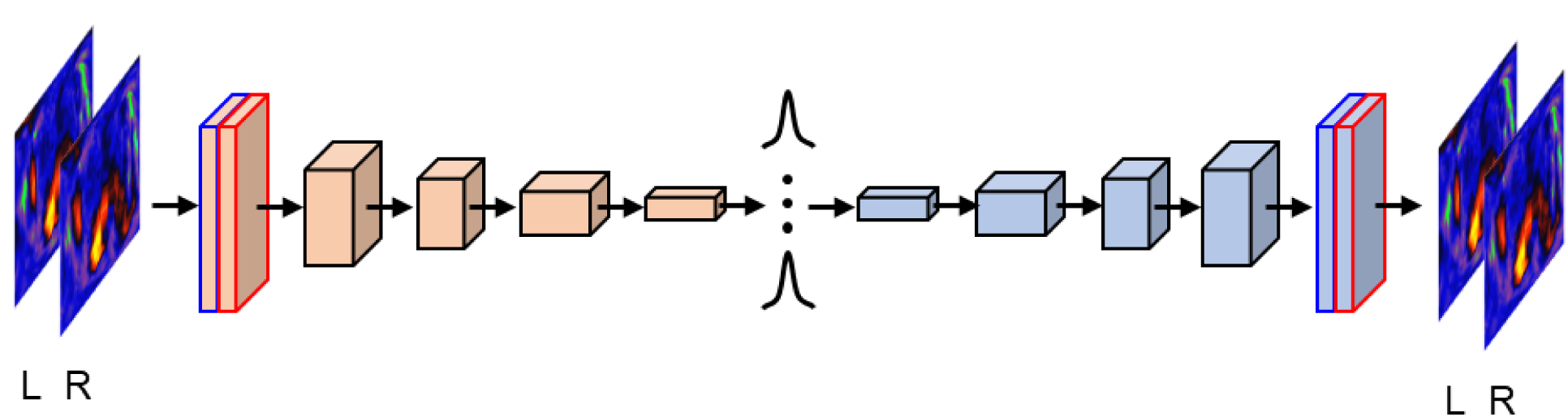
Model Architecture. Adapted from Kim, J., Zhang, Y., Han, K., Wen, Z., Choi, M., & Liu, Z. (2021). Representation learning of resting state fMRI with variational autoencoder. NeuroImage, 241, 118423. Copyright 2021 by Elsevier Inc.

#### B.2 Reconstruction performance

We demonstrate that the original FC profiles from vertices, areas, functional networks, or even functional network priors can be embedded and reconstructed using the VAE encoder and decoder, as shown in a test subject from the WU120 dataset (Supplementary Figure B.2). It is important to note that perfect reconstruction may not be ideal, as the original data could contain noise. The reconstructed data might therefore represent a denoised version of the original data.

**Supplementary Figure B.2.**
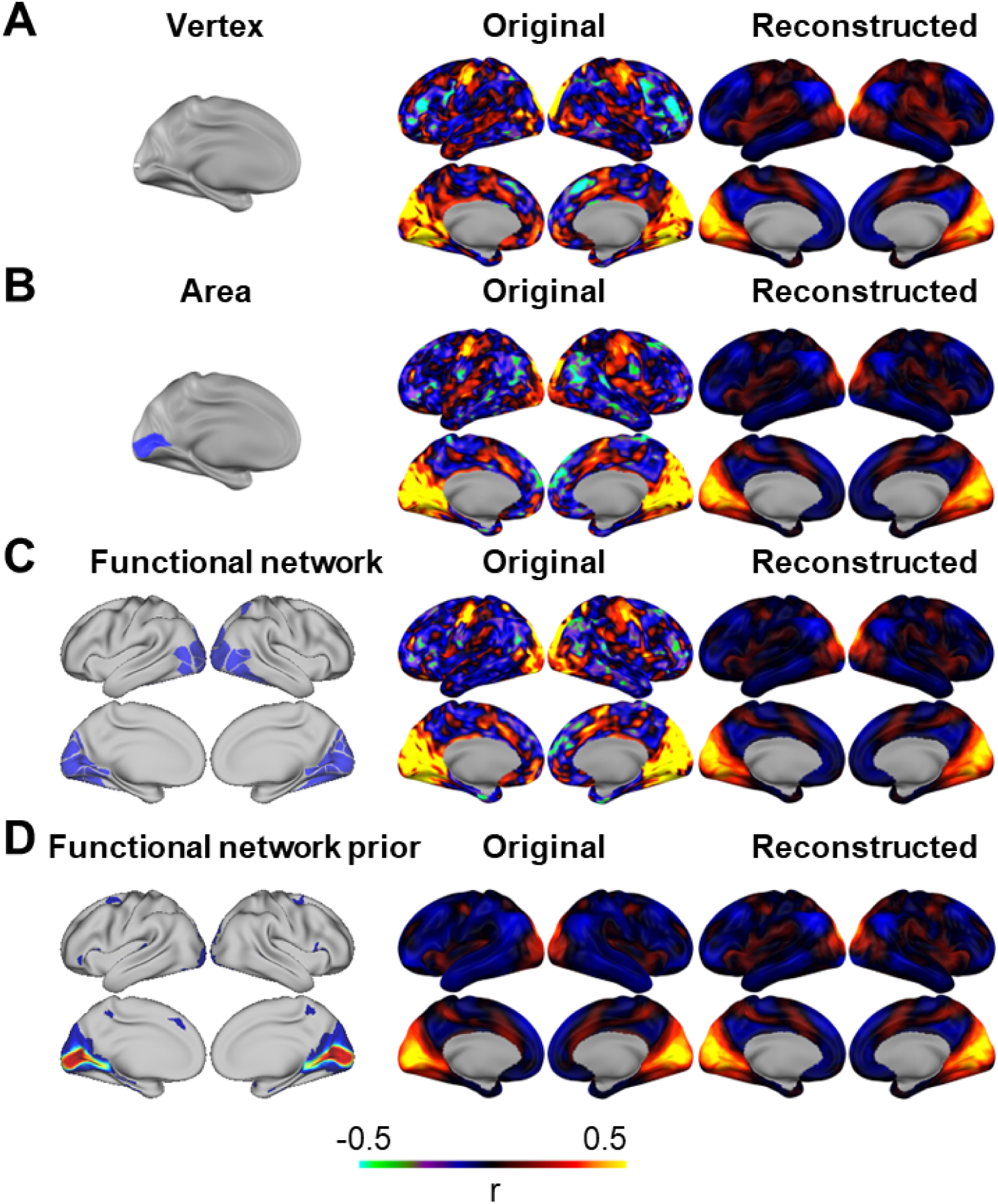
Original and reconstructed FC profiles from the beta-VAE model. A) from a vertex, B) from an area, C) from a functional network, and D) from group-averaged functional network connectivity (a.k.a. functional network priors). A-C is from one example subject in WU120. D is reproduced from data shared by Lynch, C.J., Elbau, I.G., Ng, T. et al. Frontostriatal salience network expansion in individuals in depression. Nature 633, 624–633 (2024).

#### B.3 Hyperparameter tuning

Our goal was to learn a regularized latent space that would generalize well to unseen data without significantly compromising reconstruction accuracy. To achieve this, we evaluated the reconstruction loss and KL divergence in models with β values ranging from 1 to 250. We found that β = 20 offered a good balance between reconstruction performance and KL divergence on the validation data, as shown in Supplementary Figure B.3.

**Supplementary Figure B.3.**
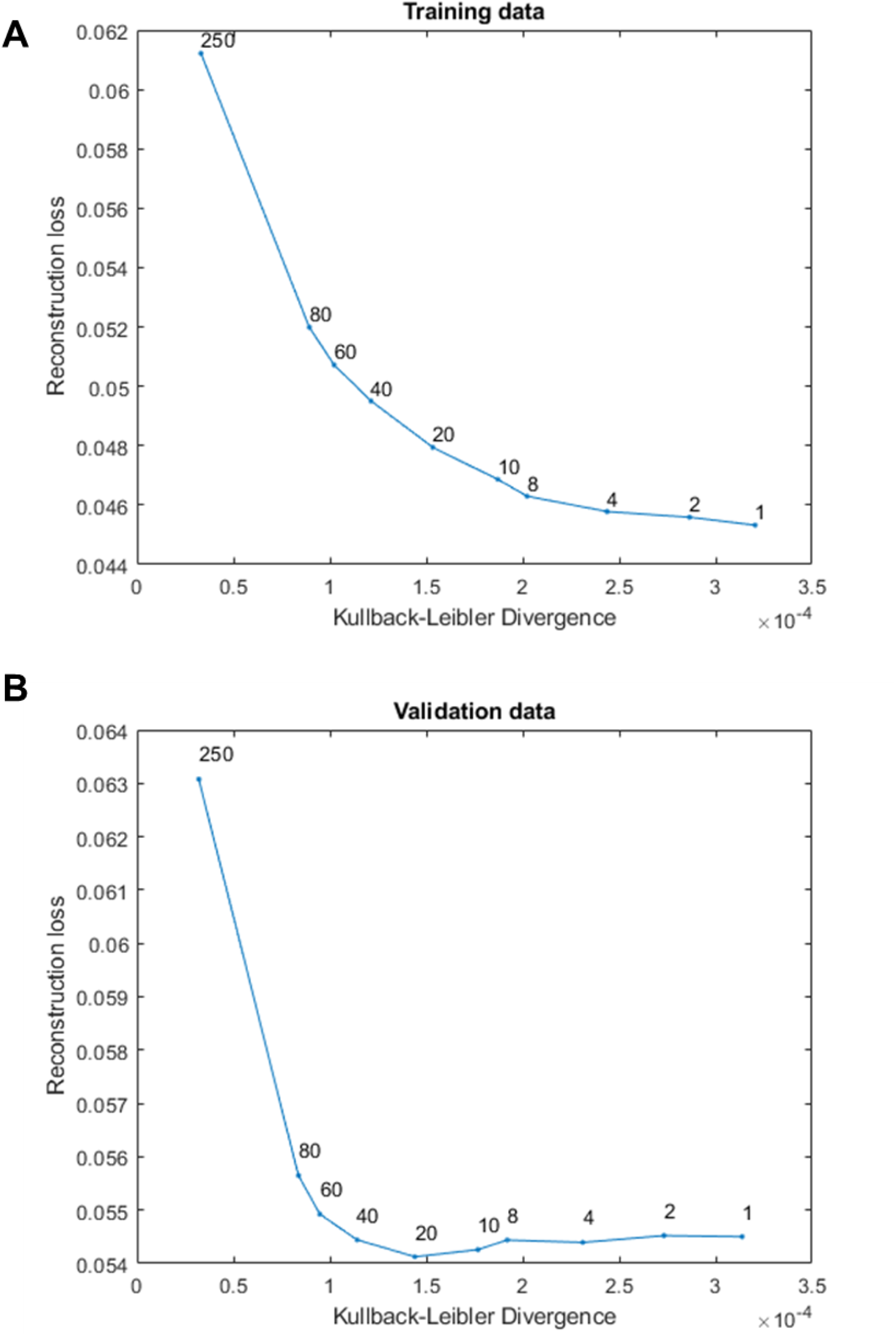
Hyperparameter tuning. A) Reconstruction loss and Kullback-Leibler divergence for different beta values in the training data. B) Reconstruction loss and Kullback-Leibler divergence for different beta values in the validation data.

### Section C. VAE Latent Space with Different Numbers of Dimensions

We observed that the mean silhouette index (SI) in the 2D latent space was significantly above zero for the average FC profile embedding from 286 areas within 12 networks across 94 HCP subjects (Figure 4). However, it was lower than the SI in the original vertex space using the correlational distance. To investigate whether certain networks would be better segregated with more latent dimensions, we replicated Figure 4 using latent dimensions of 3, 4, 8, 16, and 32. We found that both the mean SI and the correlation between distances in the latent space and vertex space increased sharply from 2 to 4 dimensions (Supplementary Figure C.1-1). The values peaked around 8 dimensions before decreasing with further increases in latent dimensions. Some networks, such as the parietal memory and fronto-parietal networks, as well as the somatomotor hand and foot networks, become more segregated at higher dimensions (Supplementary Figure C.1-2). Therefore, the 4-dimensional VAE latent space could be particularly useful for visualizing these finer separations that were not well captured in 2 dimensions (Figure 4).

**Supplementary Figure C.1-1.**
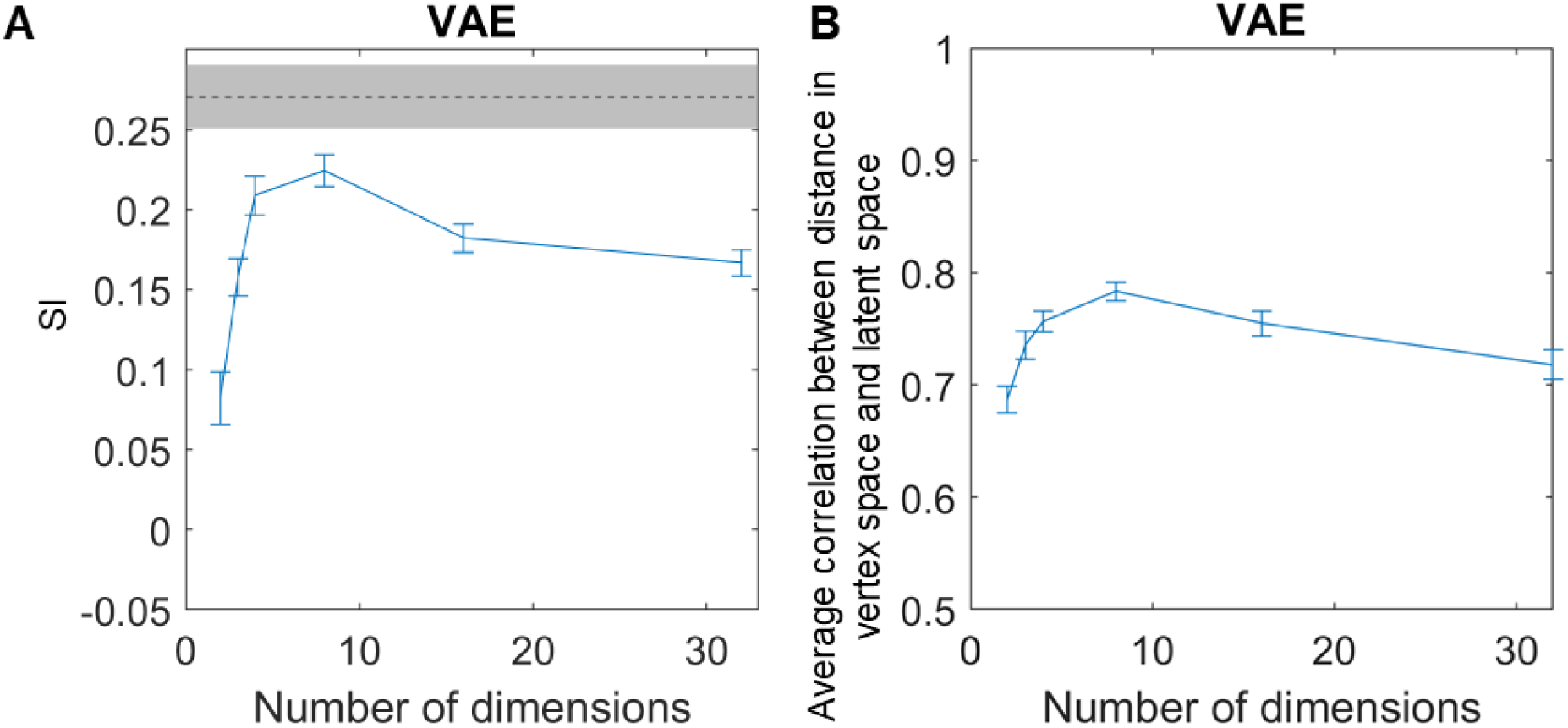
VAE Latent Space with Different Numbers of Dimensions. A) Mean SI across all areas in the 12 networks using Euclidean distance between the embeddings in the latent space. B) Average correlation between the (correlational) distance in the vertex space and the (Euclidean) distance in the latent space. The line plot shows the metric calculated with the whole 94 subject sample, and the error bars show the 95% bootstrapped confidence interval.

**Supplementary Figure C.1-2.**
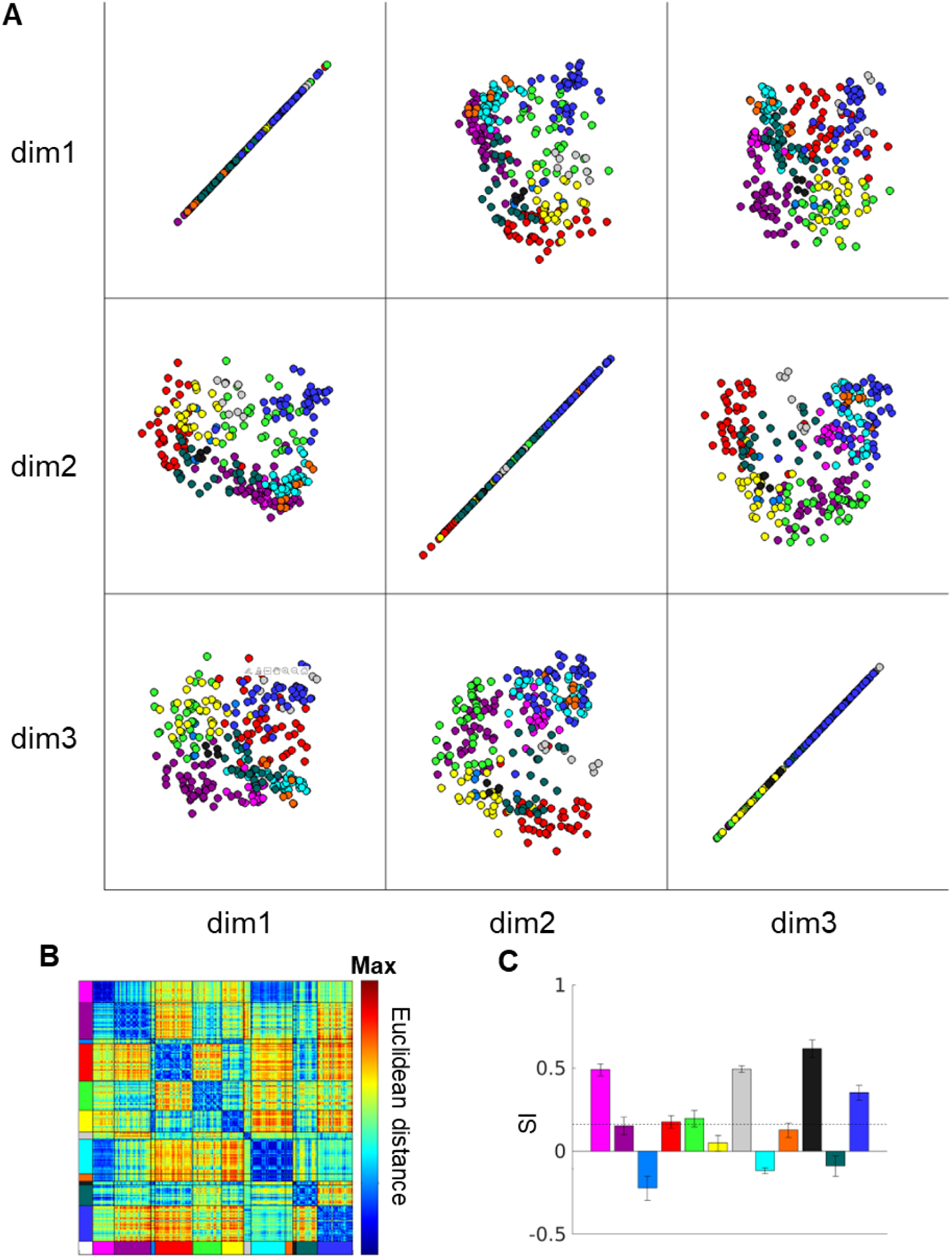
Separation of functional connectivity profiles by functional networks in the average of 94 HCP subjects (Rest1) - 3 dimensions. A) The FC profile latent embeddings with three dimensions in VAE. Each circle represents each area parcel’s mean functional connectivity profile across Rest1 sessions of 94 subjects. B) The mean Euclidean distance between the latent representations of the average across 94 subjects. C) The mean silhouette index for each functional network based on the Euclidean distance in B.

**Supplementary Figure C.1-3.**
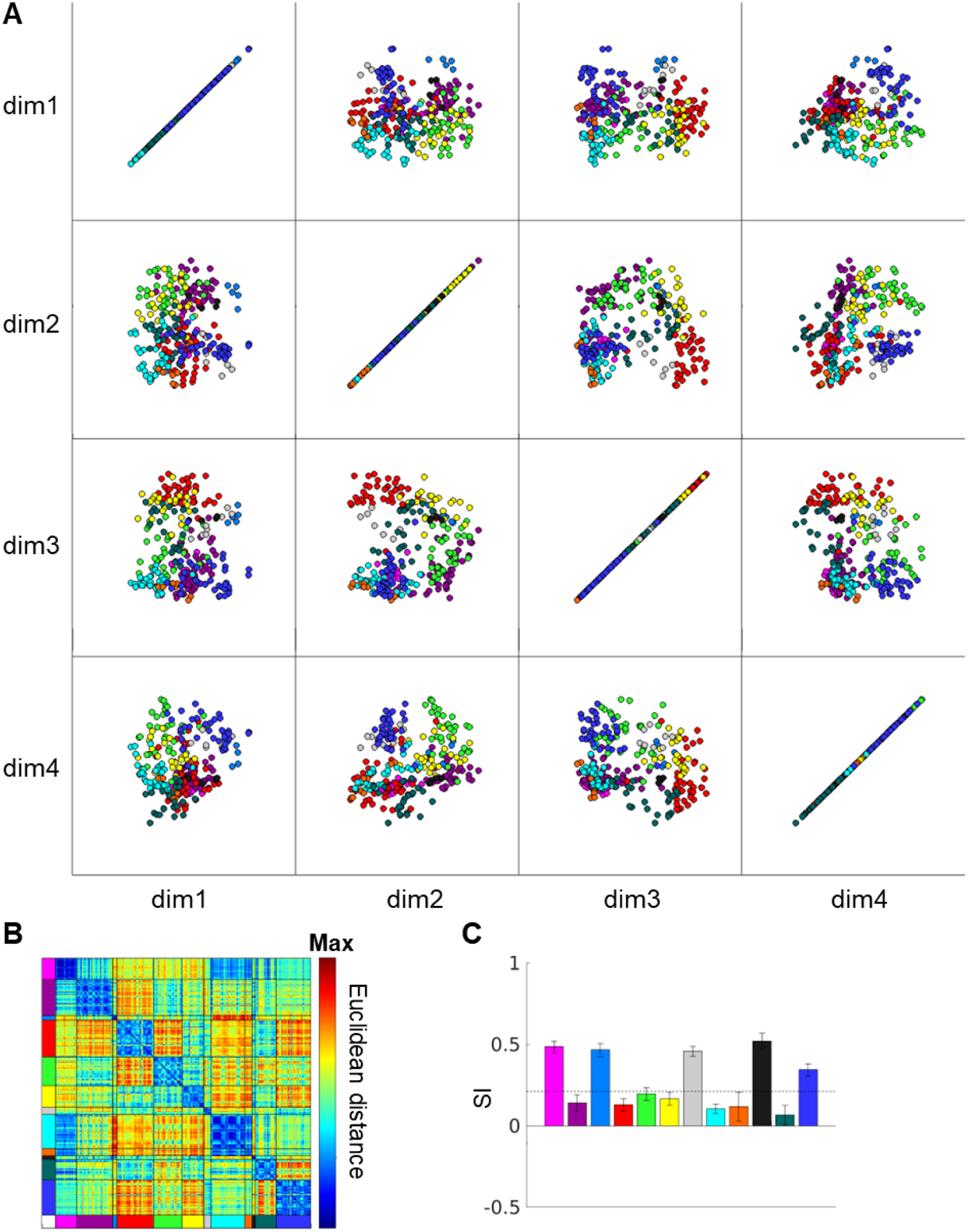
Separation of functional connectivity profiles by functional networks in the average of 94 HCP subjects (Rest1) - 4 dimensions. A) The FC profile latent embeddings with three dimensions in VAE. Each circle represents each area parcel’s mean functional connectivity profile across Rest1 sessions of 94 subjects. B) The mean Euclidean distance between the latent representations of the average across 94 subjects. C) The mean silhouette index for each functional network based on the Euclidean distance in B.

### Section D. Comparison to Alternative Autoencoder-based and Linear Dimensionality Reduction Methods

#### D.1 Alternative Autoencoder-based and Linear Dimensionality Reduction Methods

We selected the beta-VAE for its previously demonstrated ability to disentangle interpretable factors from images and to learn a continuous latent space for generating new data (Higgins et al., 2017). For the sake of completeness, we also tested the dimensionality of the FC profiles using a conventional autoencoder (Supplementary Figure D.1B), which learns a deterministic latent representation instead of a distribution, and an adversarial autoencoder (Makhzani et al., 2016) (Supplementary Figure D.1C), which employs a generative adversarial network (Goodfellow et al., 2014) based architecture to regularize the latent space.

**Figure D.1.**
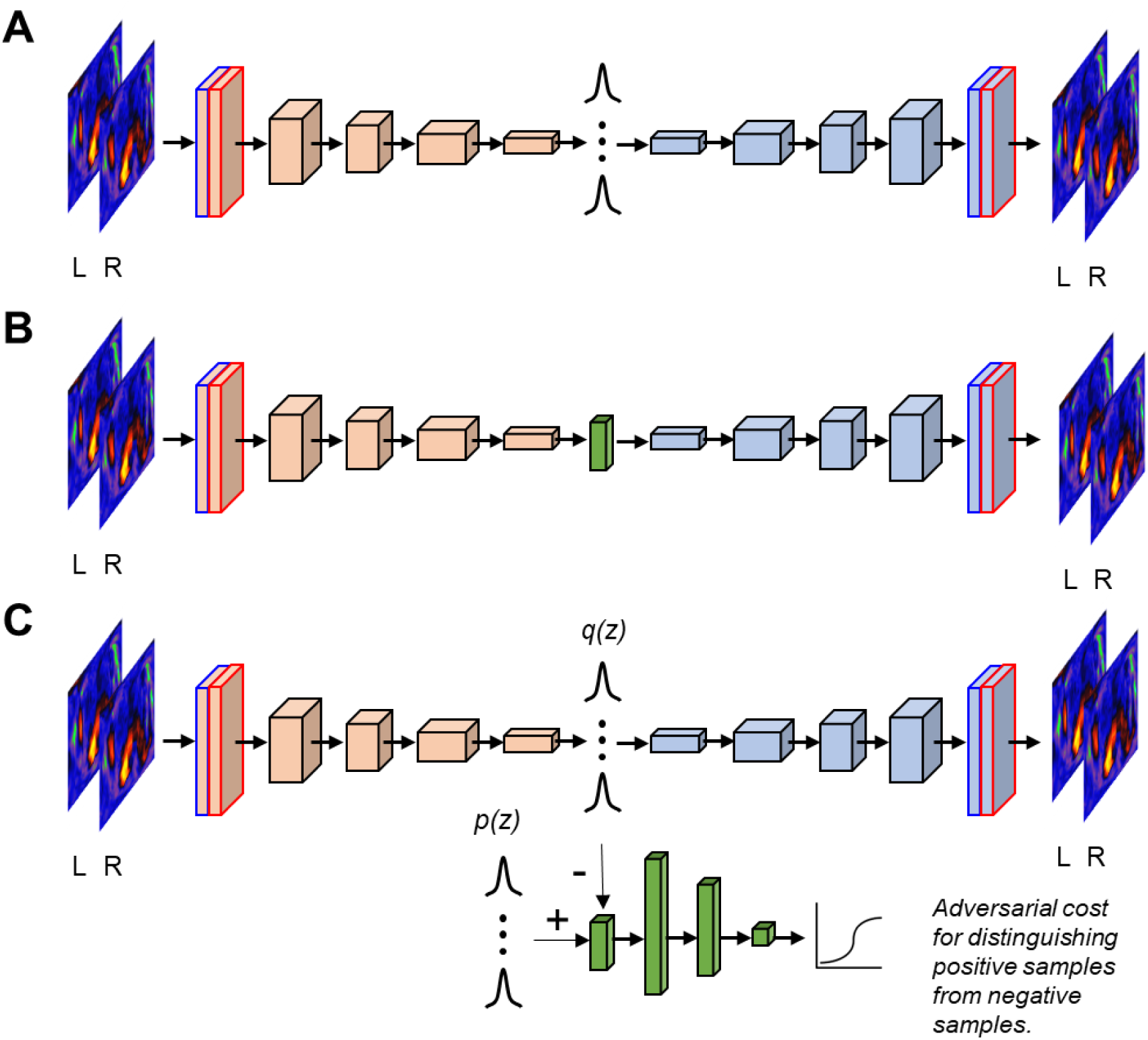
Autoencoder variants. A) Variational autoencoder. B) Autoencoder. C) Adversarial autoencoder. Blue and red boxes stand for the input images from left and right hemispheres, respectively. Adapted from Kim, J., Zhang, Y., Han, K., Wen, Z., Choi, M., \& Liu, Z. (2021). Representation learning of resting state fMRI with variational autoencoder. NeuroImage, 241, 118423. Copyright 2021 by Elsevier Inc. and Makhzani, A., Shlens, J., Jaitly, N., Goodfellow, I., Frey, B., ICLR 2016. Adversarial Autoencoders.

We chose principal component analysis (PCA) and independent component analysis (ICA) as alternative linear dimensionality reduction methods. PCA employs Singular Value Decomposition of the data, retaining only the most significant singular vectors to project the data into a lower dimensional space. ICA aims to decompose the data into a set of independent spatial maps. For memory management purposes, we performed PCA using incremental PCA (IPCA) from the Scikit-learn package (v1.3.2) in Python 3.8 (Golub & Loan, 2013; Ross et al., 2008). IPCA builds a low-rank approximation of the input data while using a constant memory amount regardless of the number of samples. ICA was then performed on the reduced data with the first 100 principal components using FastICA in Scikit-learn. To maintain fairness across algorithms, we provided the geometrically reformatted FC profile (from the 192 x 192 grid) to PCA and ICA instead of the original FC profile.

#### D.2 Single Latent Traversal

As before, we varied the magnitude of one latent dimension while keeping the other latent dimension fixed at zero. The most prominent divisions between sensorimotor networks and association networks (Margulies et al., 2016; Sydnor et al., 2021), as well as between task-positive to task-negative networks (Buckner et al., 2008; M. D. Fox et al., 2005; Raichle, 2015), remained evident in most of the autoencoder-based and linear dimensionality reduction methods. However, only the VAE models, both with regularization (β = 20) and without regularization (β = 1), were capable of demonstrating the subtle transitions across the somatomotor networks.

**Supplementary Figure D.2.**
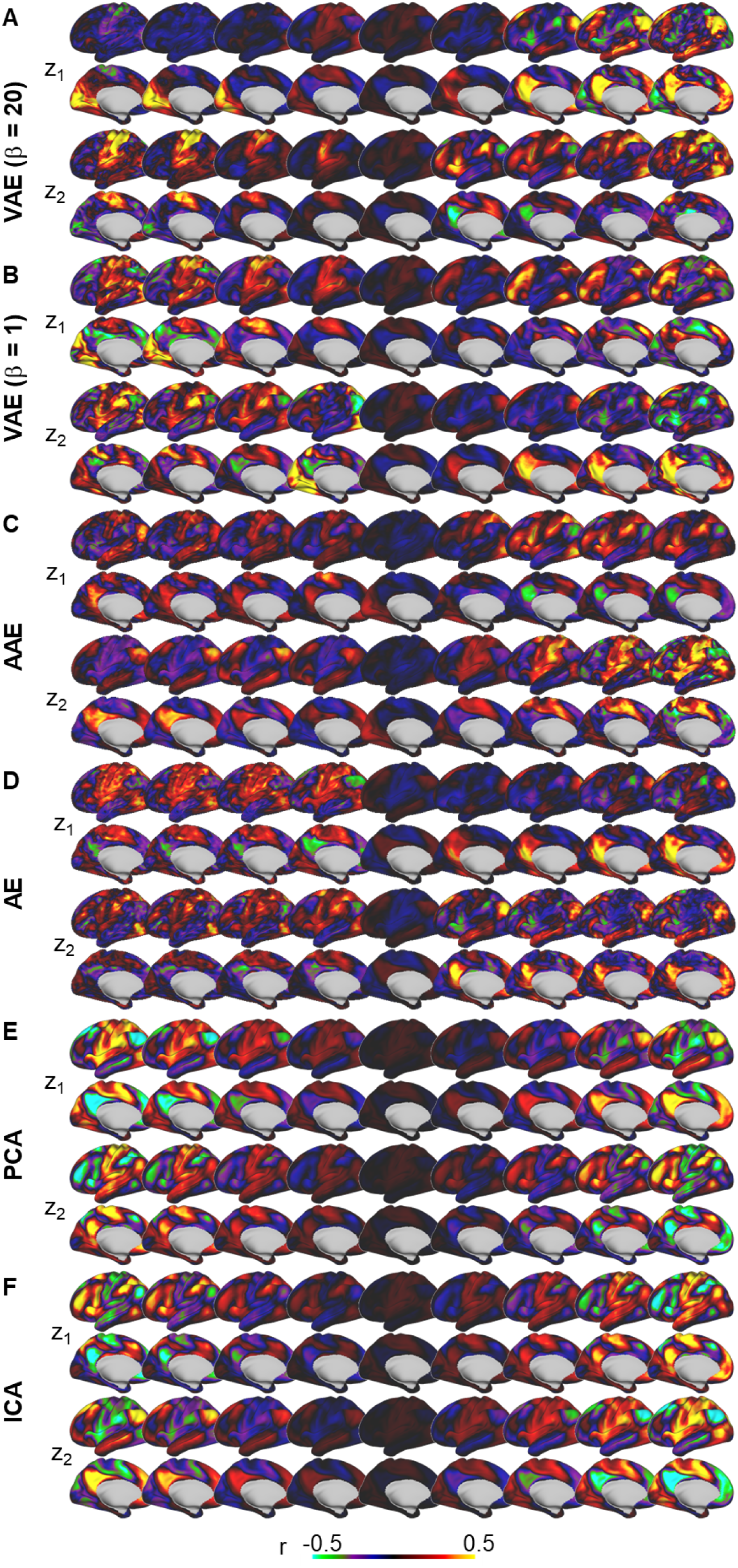
Single latent traversal. The reconstructed FC profiles from latent values of one dimension varied in equal steps from one end to the other end and the other dimension fixed to 0. A) VAE (β = 20), B) VAE (β = 1), C) AAE, D) AE, E) PCA, F) ICA.

#### D.3 Functional Networks Organization in the Latent Space

At two dimensions, the average SI for VAE (β = 20) best separated the functional networks with a mean SI of 0.086 (95% CI: [0.066, 0.099]), higher than the VAE (β = 1) (0.044, 95% CI: [0.015, 0.063]), AAE (0.035, 95% CI: [0.006,0.054]), AE (0.014, 95% CI: [-0.018,0.030]), PCA (7 × 10, 95% CI: [-0.016, 0.009]) and ICA (−0.010, 95% CI: [-0.028, 0.001]) (Supplementary Figure D.3-1 & D.3-2).

**Supplementary Figure D.3-1.**
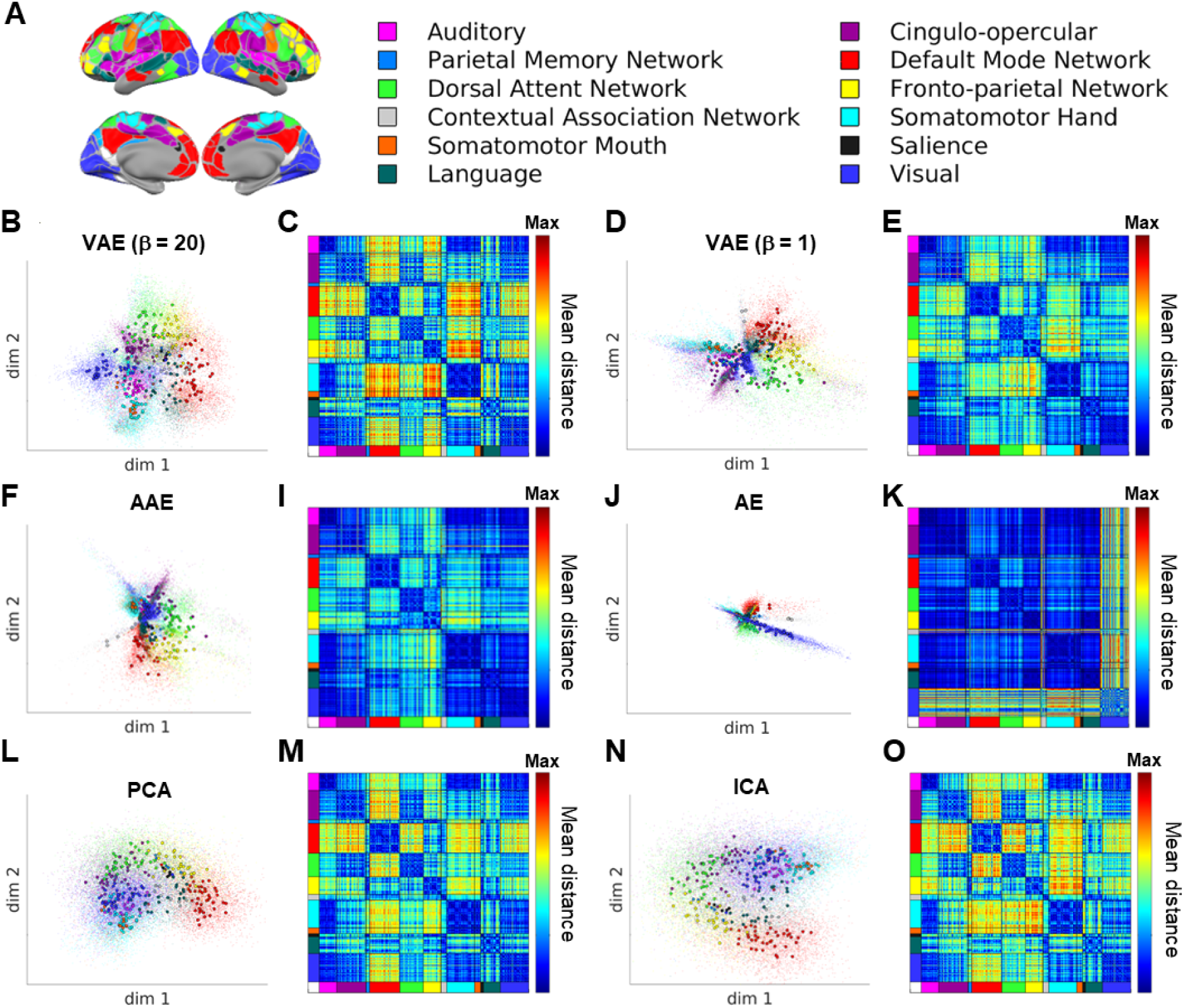
Separation of functional connectivity profiles by functional networks in the average of 94 HCP subjects (Rest1). A) Gordon network assignments for 286 area parcels. B) The FC profile latent representations with two dimensions in VAE. Each circle represents each area parcel’s mean functional connectivity profile across Rest1 sessions of 94 subjects. Each dot represents each area parcel from one subject. C) The mean Euclidean distance between the latent representations of the average across 94 subjects. D-O) Same as B-C for alternative dimensionality reduction methods. The individual dots were displayed here to demonstrate the intersubject variability in FC profile embeddings but the distance matrices (e.g. panel B) were generated using the average FC profile embeddings across subjects (circles in panel A).

**Supplementary Figure D.3-2.**
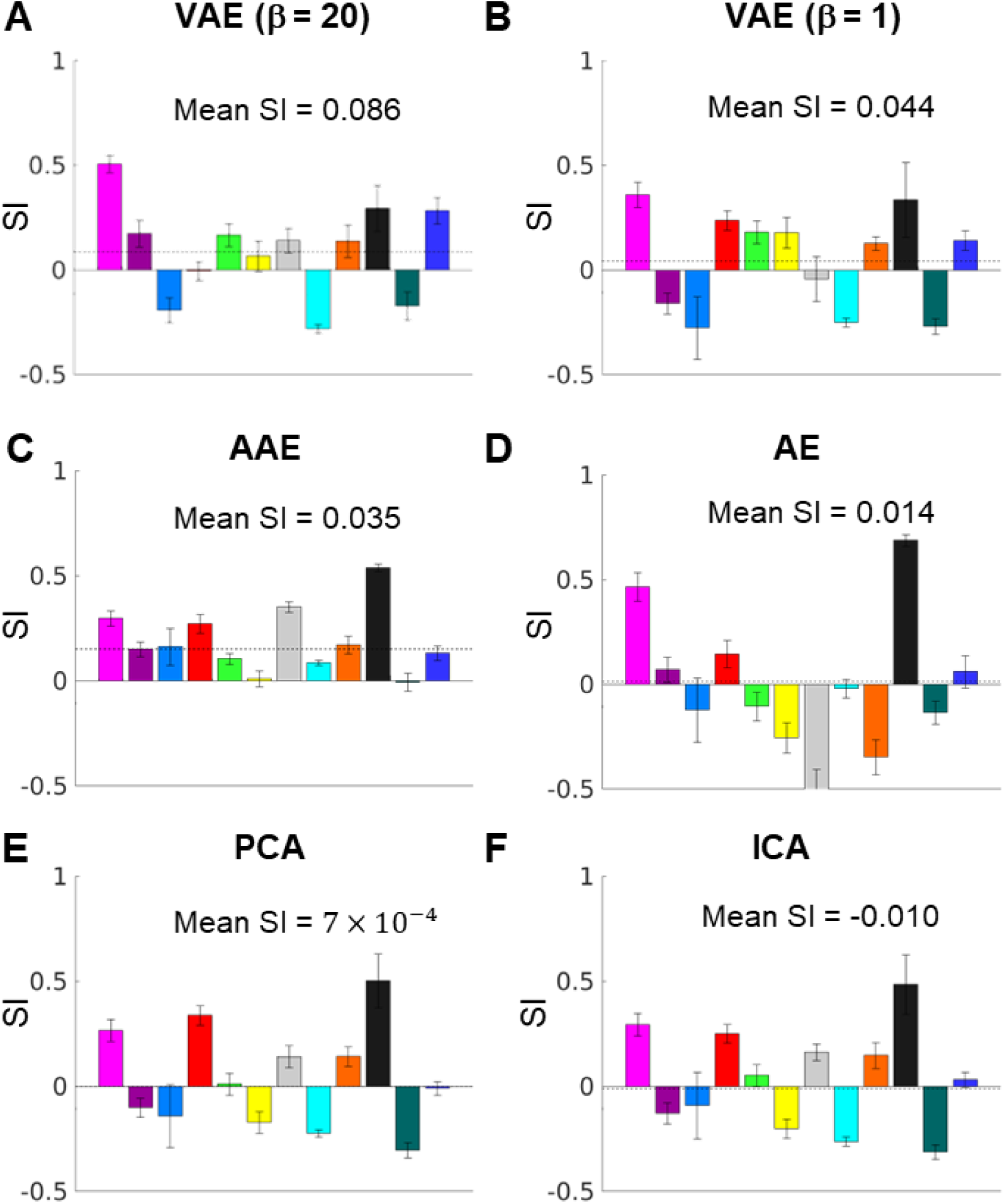
Silhouette index for each functional network based on the Euclidean distance between FC embeddings in the latent space from various dimensionality reduction methods.

#### D.4 Reconstruction Performance

Reconstruction performance was calculated with η^2^, namely, the fraction of variance in the original FC profile accounted for by variance in the reconstructed FC profile on a point-by-point basis (Cohen et al., 2008). η^2^ ranged from 0 (no similarity) to 1 (identical) and is formally defined by:

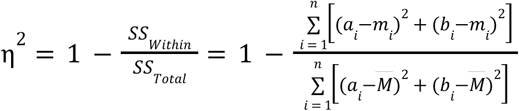

where *a*_*i*_ and *b*_*i*_ represent the values at position *i* in maps *a* and *b*, respectively. *m*_*i*_ is the mean value of the two images at position 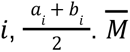 is the grand mean value across the mean image *m*. Self-connectivity at all positions was excluded. This similarity matrix is sensitive to the difference in *a* and *b* in scales and offsets. For convenience, we used the inversely formatted FC profiles from the 2D image in Figure 1A as the reference ground truth data (η^2^ > 0.99) to the original data) to be compared with the reconstructed data from latent representations. We computed the reconstruction performance on the FC profiles from each of the 333 area parcels (Gordon et al., 2016) in the 10 test subjects in the WU120 dataset and the two sessions (Rest1 and Rest2) for the 94 test subjects in the HCP dataset to test the out-of-sample and out-of-distribution generalization, respectively. A good latent representation should reflect a trait-like property of the subject and minimize across session variation, yet distinguishable from other subjects. Since the HCP data had two sessions, we calculated two additional reference measure: 1) the η^2^ between the original data in HCP Rest1 and HCP Rest2 of the same subject, which provides the noise ceiling of the reconstruction, and 2) the η^2^ between each subject to the rest 93 subjects in each session (and then averaged across the Rest1 and Rest2 sessions), which provides the mean baseline.

In the reconstructed FC profile from one example area parcel (parcel 15 in the medial visual cortex, Supplementary Figure D.4-1, a general trend for more detailed variations in reconstruction FC was observed with an increasing number of dimensions (Supplementary Figure D.4-1A). Overall, the mean reconstruction performance across all parcels in each individual was similar across methods. Autoencoder-based latent representations provided better reconstruction performance when the latent representation had only 2 dimensions (Supplementary Figure D.4-1B) while linear methods provided marginally better reconstruction performance at 32 dimensions (Supplementary Figure D.4-1D). In all cases, the reconstruction performance from the latent representations was, on average higher than the mean baseline, suggesting that the latent representations captured individual-specific features in additional to group-average features. With 32 dimensions, the reconstruction performance was approaching the noise ceiling Supplementary Figure D4-1D). This observation persisted when each individual subject’s average reconstruction performance was normalized to their own noise ceiling (Supplementary Figure D.4-2).

**Supplementary Figure D.4-1.**
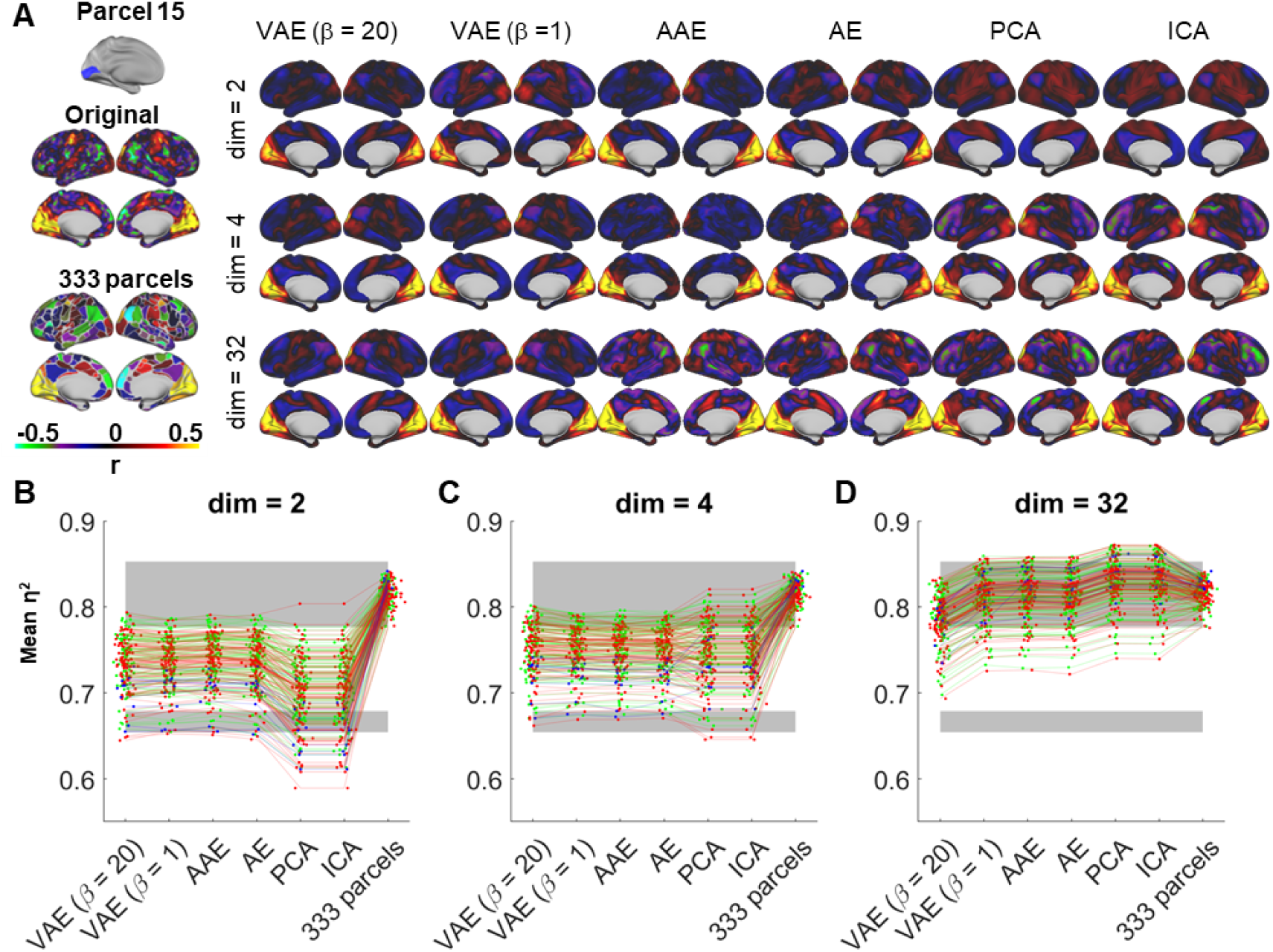
Reconstruction performance comparison across dimensionality reduction methods. A) Visualization of the original and reconstructed FC profiles from parcel 15 in an example subject (subject 111). B-D) The reconstruction performance for all subjects in the HCP and WU120 test datasets. Green line: HCP Rest1, Red line: HCP Rest2, Blue line: WU120. Gray shaded area: mean and standard deviation of the noise ceiling and the mean baseline across 94 HCP subjects. N.B.: 333 parcels always have 333 dimensions and were repeatedly displayed in all three panels as a reference. VAE = variational autoencoder. AAE = adversarial autoencoder. AE = autoencoder. PCA = principal component analysis. ICA = independent component analysis.

**Supplementary Figure D.4-2.**
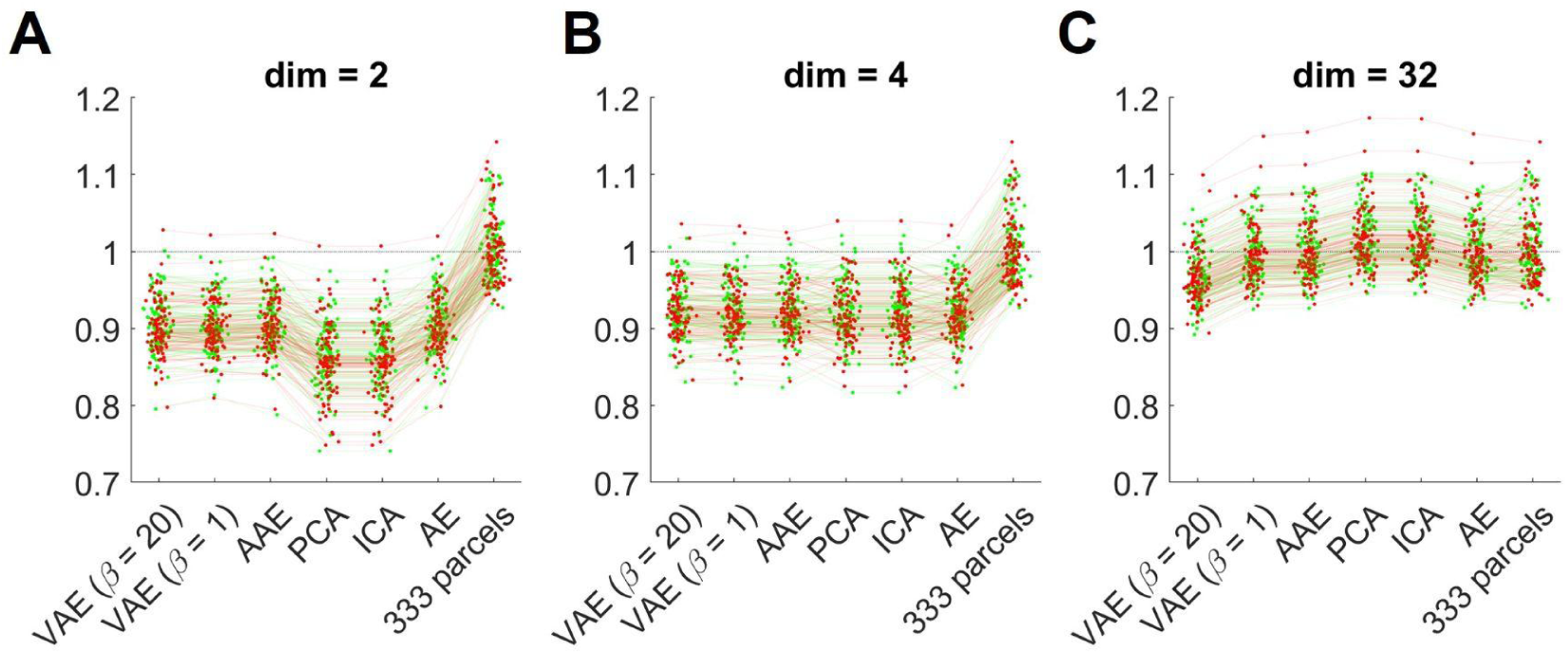
Reconstruction performance of the HCP sessions (188 sessions across 94 subjects) normalized to the noise ceiling. Green line: HCP Rest1, Red line: HCP Rest2, Blue line: WU120. Gray shaded area: mean and standard deviation of the noise ceiling and the mean baseline. N.B.: 333 parcels always have 333 dimensions and were repeatedly displayed in all three panels as a reference. VAE = variational autoencoder. AAE = adversarial autoencoder. AE = autoencoder. PCA = principal component analysis. ICA = independent component analysis.

## Reference

Akiki, T. J., & Abdallah, C. G. (2019). Determining the Hierarchical Architecture of the Human Brain Using Subject-Level Clustering of Functional Networks. Scientific Reports, 9(1), Article 1. 10.1038/s41598-019-55738-y

Allen, E. J., St-Yves, G., Wu, Y., Breedlove, J. L., Prince, J. S., Dowdle, L. T., Nau, M., Caron, B., Pestilli, F., Charest, I., Hutchinson, J. B., Naselaris, T., & Kay, K. (2022). A massive 7T fMRI dataset to bridge cognitive neuroscience and artificial intelligence. Nature Neuroscience, 25(1), 116–126. 10.1038/s41593-021-00962-x

Andrews-Hanna, J. R., Reidler, J. S., Sepulcre, J., Poulin, R., & Buckner, R. L. (2010). Functional-Anatomic Fractionation of the Brain’s Default Network. Neuron, 65(4). 10.1016/j.neuron.2010.02.005

Arslan, S., Ktena, S. I., Makropoulos, A., Robinson, E. C., Rueckert, D., & Parisot, S. (2018). Human brain mapping: A systematic comparison of parcellation methods for the human cerebral cortex. NeuroImage, 170, 5–30. 10.1016/j.neuroimage.2017.04.014

Atasoy, S., Deco, G., Kringelbach, M. L., & Pearson, J. (2018). Harmonic Brain Modes: A Unifying Framework for Linking Space and Time in Brain Dynamics. The Neuroscientist, 24(3), 277–293. 10.1177/1073858417728032

Atasoy, S., Donnelly, I., & Pearson, J. (2016). Human brain networks function in connectome-specific harmonic waves. Nature Communications, 7(1), 10340. 10.1038/ncomms10340

Bassett, D. S., Wymbs, N. F., Porter, M. A., Mucha, P. J., Carlson, J. M., & Grafton, S. T. (2011). Dynamic reconfiguration of human brain networks during learning. Proceedings of the National Academy of Sciences, 108(18), 7641–7646. 10.1073/pnas.1018985108

Betzel, R. F., Bertolero, M. A., Gordon, E. M., Gratton, C., Dosenbach, N. U. F., & Bassett, D. S. (2019). The community structure of functional brain networks exhibits scale-specific patterns of inter- and intra-subject variability. NeuroImage, 202. 10.1016/j.neuroimage.2019.07.003

Betzel, R. F., Byrge, L., He, Y., Goñi, J., Zuo, X.-N., & Sporns, O. (2014). Changes in structural and functional connectivity among resting-state networks across the human lifespan. NeuroImage, 102, 345–357. 10.1016/j.neuroimage.2014.07.067

Bijsterbosch, J. D., Woolrich, M. W., Glasser, M. F., Robinson, E. C., Beckmann, C. F., Van Essen, D. C., Harrison, S. J., & Smith, S. M. (2018). The relationship between spatial configuration and functional connectivity of brain regions. eLife, 7, e32992. 10.7554/eLife.32992

Bookstein, F. L. (1997). Two shape metrics for biomedical outline data: Bending energy, Procrustes distance, and the biometrical modeling of shape phenomena. Proceedings of 1997 International Conference on Shape Modeling and Applications, 110–120. 10.1109/SMA.1997.634888

Braga, R. M., & Buckner, R. L. (2017). Parallel Interdigitated Distributed Networks within the Individual Estimated by Intrinsic Functional Connectivity. Neuron, 95(2), 457–471.e5. 10.1016/j.neuron.2017.06.038

Brett, M., Johnsrude, I. S., & Owen, A. M. (2002). The problem of functional localization in the human brain. Nature Reviews Neuroscience, 3(3), 243–249. 10.1038/nrn756

Buckner, R. L., Andrews-Hanna, J. R., & Schacter, D. L. (2008). The Brain’s Default Network. Annals of the New York Academy of Sciences, 1124(1), 1–38. 10.1196/annals.1440.011

Buckner, R. L., Krienen, F. M., & Yeo, B. T. T. (2013). Opportunities and limitations of intrinsic functional connectivity MRI. Nature Neuroscience, 16(7), 832–837. 10.1038/nn.3423

Calhoun, V. d., Adali, T., Pearlson, G. d., & Pekar, J. j. (2001). A method for making group inferences from functional MRI data using independent component analysis. Human Brain Mapping, 14(3), 140–151. 10.1002/hbm.1048

Churchland, M. M., Cunningham, J. P., Kaufman, M. T., Foster, J. D., Nuyujukian, P., Ryu, S. I., & Shenoy, K. V. (2012). Neural population dynamics during reaching. Nature, 487(7405), 51–56. 10.1038/nature11129

Cohen, A. L., Fair, D. A., Dosenbach, N. U. F., Miezin, F. M., Dierker, D., Van Essen, D. C., Schlaggar, B. L., & Petersen, S. E. (2008). Defining functional areas in individual human brains using resting functional connectivity MRI. NeuroImage, 41(1), 45–57. 10.1016/j.neuroimage.2008.01.066

Cole, M. W., Ito, T., Bassett, D. S., & Schultz, D. H. (2016). Activity flow over resting-state networks shapes cognitive task activations. Nature Neuroscience, 19(12), Article 12. 10.1038/nn.4406

Craddock, R. C., James, G. A., Holtzheimer III, P. E., Hu, X. P., & Mayberg, H. S. (2012). A whole brain fMRI atlas generated via spatially constrained spectral clustering. Human Brain Mapping, 33(8), 1914–1928. 10.1002/hbm.21333

Cui, Z., Li, H., Xia, C. H., Larsen, B., Adebimpe, A., Baum, G. L., Cieslak, M., Gur, R. E., Gur, R. C., Moore, T. M., Oathes, D. J., Alexander-Bloch, A. F., Raznahan, A., Roalf, D. R., Shinohara, R. T., Wolf, D. H., Davatzikos, C., Bassett, D. S., Fair, D. A., … Satterthwaite, T. D. (2020). Individual Variation in Functional Topography of Association Networks in Youth. Neuron, 106(2), 340–353.e8. 10.1016/j.neuron.2020.01.029

Cusack, R., Brett, M., & Osswald, K. (2003). An Evaluation of the Use of Magnetic Field Maps to Undistort Echo-Planar Images. NeuroImage, 18(1), 127–142. 10.1006/nimg.2002.1281

Damoiseaux, J. S., Rombouts, S. a. R. B., Barkhof, F., Scheltens, P., Stam, C. J., Smith, S. M., & Beckmann, C. F. (2006). Consistent resting-state networks across healthy subjects. Proceedings of the National Academy of Sciences, 103(37), 13848–13853. 10.1073/pnas.0601417103

Dong, H.-M., Margulies, D. S., Zuo, X.-N., & Holmes, A. J. (2021). Shifting gradients of macroscale cortical organization mark the transition from childhood to adolescence. Proceedings of the National Academy of Sciences, 118(28), e2024448118. 10.1073/pnas.2024448118

Dworetsky, A., Seitzman, B. A., Adeyemo, B., Neta, M., Coalson, R. S., Petersen, S. E., & Gratton, C. (2021). Probabilistic mapping of human functional brain networks identifies regions of high group consensus. NeuroImage, 237, 118164. 10.1016/j.neuroimage.2021.118164

Dworetsky, A., Seitzman, B. A., Adeyemo, B., Nielsen, A. N., Hatoum, A. S., Smith, D. M., Nichols, T. E., Neta, M., Petersen, S. E., & Gratton, C. (2024). Two common and distinct forms of variation in human functional brain networks. Nature Neuroscience, 27(6), 1187–1198. 10.1038/s41593-024-01618-2

Fair, D. A. (2020). Correction of respiratory artifacts in MRI head motion estimates. NeuroImage, 17.

Feczko, E., Conan, G., Marek, S., Tervo-Clemmens, B., Cordova, M., Doyle, O., Earl, E., Perrone, A., Sturgeon, D., Klein, R., Harman, G., Kilamovich, D., Hermosillo, R., Miranda-Dominguez, O., Adebimpe, A., Bertolero, M., Cieslak, M., Covitz, S., Hendrickson, T., … Fair, D. A. (2021). Adolescent Brain Cognitive Development (ABCD) Community MRI Collection and Utilities (p. 2021.07.09.451638). bioRxiv. 10.1101/2021.07.09.451638

Fischl, B. (2012). FreeSurfer. NeuroImage, 62(2), 774–781. 10.1016/j.neuroimage.2012.01.021

Fox, M. D., Liu, H., & Pascual-Leone, A. (2013). Identification of reproducible individualized targets for treatment of depression with TMS based on intrinsic connectivity. NeuroImage, 66, 151–160. 10.1016/j.neuroimage.2012.10.082

Fox, M. D., Snyder, A. Z., Vincent, J. L., Corbetta, M., Van Essen, D. C., & Raichle, M. E. (2005). The human brain is intrinsically organized into dynamic, anticorrelated functional networks. Proceedings of the National Academy of Sciences, 102(27), 9673–9678. 10.1073/pnas.0504136102

Fox, P. T., & Friston, K. J. (2012). Distributed processing; distributed functions? NeuroImage, 61(2), 407–426. 10.1016/j.neuroimage.2011.12.051

Gao, W., Alcauter, S., Elton, A., Hernandez-Castillo, C. R., Smith, J. K., Ramirez, J., & Lin, W. (2015). Functional Network Development During the First Year: Relative Sequence and Socioeconomic Correlations. Cerebral Cortex (New York, N.Y.: 1991), 25(9), 2919–2928. 10.1093/cercor/bhu088

Glasser, M. F., Coalson, T. S., Robinson, E. C., Hacker, C. D., Harwell, J., Yacoub, E., Ugurbil, K., Andersson, J., Beckmann, C. F., Jenkinson, M., Smith, S. M., & Van Essen, D. C. (2016). A multi-modal parcellation of human cerebral cortex. Nature, 536(7615), 171–178. 10.1038/nature18933

Glasser, M. F., Sotiropoulos, S. N., Wilson, J. A., Coalson, T. S., Fischl, B., Andersson, J. L., Xu, J., Jbabdi, S., Webster, M., Polimeni, J. R., Van Essen, D. C., & Jenkinson, M. (2013). The minimal preprocessing pipelines for the Human Connectome Project. NeuroImage, 80, 105–124. 10.1016/j.neuroimage.2013.04.127

Glasser, M. F., & Van Essen, D. C. (2011). Mapping Human Cortical Areas In Vivo Based on Myelin Content as Revealed by T1- and T2-Weighted MRI. Journal of Neuroscience, 31(32), 11597–11616. 10.1523/JNEUROSCI.2180-11.2011

Golub, G. H., & Loan, C. F. V. (2013). Matrix Computations. JHU Press.

Goodfellow, I., Pouget-Abadie, J., Mirza, M., Xu, B., Warde-Farley, D., Ozair, S., Courville, A., & Bengio, Y. (2014). Generative Adversarial Nets. Advances in Neural Information Processing Systems, 27. https://papers.nips.cc/paper_files/paper/2014/hash/5ca3e9b122f61f8f06494c97b1afccf3-Abstract.html

Gordon, E. M., Laumann, T. O., Adeyemo, B., Gilmore, A. W., Nelson, S. M., Dosenbach, N. U. F., & Petersen, S. E. (2017). Individual-specific features of brain systems identified with resting state functional correlations. NeuroImage, 146, 918–939. 10.1016/j.neuroimage.2016.08.032

Gordon, E. M., Laumann, T. O., Adeyemo, B., Huckins, J. F., Kelley, W. M., & Petersen, S. E. (2016). Generation and Evaluation of a Cortical Area Parcellation from Resting-State Correlations. Cerebral Cortex, 26(1), 288–303. 10.1093/cercor/bhu239

Gordon, E. M., Laumann, T. O., Adeyemo, B., & Petersen, S. E. (2017). Individual Variability of the System-Level Organization of the Human Brain. Cerebral Cortex, 27(1), 386–399. 10.1093/cercor/bhv239

Gordon, E. M., Laumann, T. O., Gilmore, A. W., Newbold, D. J., Greene, D. J., Berg, J. J., Ortega, M., Hoyt-Drazen, C., Gratton, C., Sun, H., Hampton, J. M., Coalson, R. S., Nguyen, A. L., McDermott, K. B., Shimony, J. S., Snyder, A. Z., Schlaggar, B. L., Petersen, S. E., Nelson, S. M., & Dosenbach, N. U. F. (2017). Precision Functional Mapping of Individual Human Brains. Neuron, 95(4). 10.1016/j.neuron.2017.07.011

Gordon, E. M., Laumann, T. O., Marek, S., Raut, R. V., Gratton, C., Newbold, D. J., Greene, D. J., Coalson, R. S., Snyder, A. Z., Schlaggar, B. L., Petersen, S. E., Dosenbach, N. U. F., & Nelson, S. M. (2020). Default-mode network streams for coupling to language and control systems. Proceedings of the National Academy of Sciences of the United States of America, 117(29), 17308–17319. 10.1073/pnas.2005238117

Gotts, S. J., Gilmore, A. W., & Martin, A. (2020). Brain networks, dimensionality, and global signal averaging in resting-state fMRI: Hierarchical network structure results in low-dimensional spatiotemporal dynamics. NeuroImage, 205, 116289. 10.1016/j.neuroimage.2019.116289

Gratton, C., Dworetsky, A., Coalson, R. S., Adeyemo, B., Laumann, T. O., Wig, G. S., Kong, T. S., Gratton, G., Fabiani, M., Barch, D. M., Tranel, D., Miranda-Dominguez, O., Fair, D. A., Dosenbach, N. U. F., Snyder, A. Z., Perlmutter, J. S., Petersen, S. E., & Campbell, M. C. (2020). Removal of high frequency contamination from motion estimates in single-band fMRI saves data without biasing functional connectivity. NeuroImage, 217, 116866. 10.1016/j.neuroimage.2020.116866

Gratton, C., Kraus, B. T., Greene, D. J., Gordon, E. M., Laumann, T. O., Nelson, S. M., Dosenbach, N. U. F., & Petersen, S. E. (2020). Defining Individual-Specific Functional Neuroanatomy for Precision Psychiatry. Biological Psychiatry, 88(1), 28–39. 10.1016/j.biopsych.2019.10.026

Gratton, C., Laumann, T. O., Nielsen, A. N., Greene, D. J., Gordon, E. M., Gilmore, A. W., Nelson, S. M., Coalson, R. S., Snyder, A. Z., Schlaggar, B. L., Dosenbach, N. U. F., & Petersen, S. E. (2018). Functional Brain Networks Are Dominated by Stable Group and Individual Factors, Not Cognitive or Daily Variation. Neuron, 98(2), 439–452.e5. 10.1016/j.neuron.2018.03.035

Grayson, D. S., & Fair, D. A. (2017). Development of large-scale functional networks from birth to adulthood: A guide to the neuroimaging literature. NeuroImage, 160, 15–31. 10.1016/j.neuroimage.2017.01.079

Guntupalli, J. S., Feilong, M., & Haxby, J. V. (2018). A computational model of shared fine-scale structure in the human connectome. PLOS Computational Biology, 14(4), e1006120. 10.1371/journal.pcbi.1006120

Haak, K. V., Marquand, A. F., & Beckmann, C. F. (2018). Connectopic mapping with resting-state fMRI. NeuroImage, 170, 83–94. 10.1016/j.neuroimage.2017.06.075

Hacker, C. D., Laumann, T. O., Szrama, N. P., Baldassarre, A., Snyder, A. Z., Leuthardt, E. C., & Corbetta, M. (2013). Resting state network estimation in individual subjects. NeuroImage, 82, 616–633. 10.1016/j.neuroimage.2013.05.108

Han, L., Savalia, N. K., Chan, M. Y., Agres, P. F., Nair, A. S., & Wig, G. S. (2018). Functional Parcellation of the Cerebral Cortex Across the Human Adult Lifespan. Cerebral Cortex, 28(12), 4403–4423. 10.1093/cercor/bhy218

Haxby, J. V., Guntupalli, J. S., Nastase, S. A., & Feilong, M. (2020). Hyperalignment: Modeling shared information encoded in idiosyncratic cortical topographies. eLife, 9, e56601. 10.7554/eLife.56601

Hermosillo, R. J. M., Moore, L. A., Feczko, E., Miranda-Domínguez, Ó., Pines, A., Dworetsky, A., Conan, G., Mooney, M. A., Randolph, A., Graham, A., Adeyemo, B., Earl, E., Perrone, A., Carrasco, C. M., Uriarte-Lopez, J., Snider, K., Doyle, O., Cordova, M., Koirala, S., … Fair, D. A. (2024). A precision functional atlas of personalized network topography and probabilities. Nature Neuroscience, 27(5), 1000–1013. 10.1038/s41593-024-01596-5

Higgins, I., Matthey, L., Pal, A., Burgess, C., Glorot, X., Botvinick, M., Mohamed, S., & Lerchner, A. (2017). β-VAE: LEARNING BASIC VISUAL CONCEPTS WITH A CONSTRAINED VARIATIONAL FRAMEWORK. ICLR.

Hong, S.-J., Vos de Wael, R., Bethlehem, R. A. I., Lariviere, S., Paquola, C., Valk, S. L., Milham, M. P., Di Martino, A., Margulies, D. S., Smallwood, J., & Bernhardt, B. C. (2019). Atypical functional connectome hierarchy in autism. Nature Communications, 10(1), Article 1. 10.1038/s41467-019-08944-1

Howell, B. R., Styner, M. A., Gao, W., Yap, P.-T., Wang, L., Baluyot, K., Yacoub, E., Chen, G., Potts, T., Salzwedel, A., Li, G., Gilmore, J. H., Piven, J., Smith, J. K., Shen, D., Ugurbil, K., Zhu, H., Lin, W., & Elison, J. T. (2019). The UNC/UMN Baby Connectome Project (BCP): An overview of the study design and protocol development. NeuroImage, 185, 891–905. 10.1016/j.neuroimage.2018.03.049

Jezzard, P., & Balaban, R. S. (1995). Correction for geometric distortion in echo planar images from B0 field variations. Magnetic Resonance in Medicine, 34(1), 65–73.

Kaplan, S., Meyer, D., Miranda-Dominguez, O., Perrone, A., Earl, E., Alexopoulos, D., Barch, D. M., Day, T. K. M., Dust, J., Eggebrecht, A. T., Feczko, E., Kardan, O., Kenley, J. K., Rogers, C. E., Wheelock, M. D., Yacoub, E., Rosenberg, M., Elison, J. T., Fair, D. A., & Smyser, C. D. (2022). Filtering respiratory motion artifact from resting state fMRI data in infant and toddler populations. NeuroImage, 247, 118838. 10.1016/j.neuroimage.2021.118838

Kim, J.-H., De Asis-Cruz, J., Krishnamurthy, D., & Limperopoulos, C. (2023). Toward a more informative representation of the fetal–neonatal brain connectome using variational autoencoder. eLife, 12, e80878. 10.7554/eLife.80878

Kim, J.-H., De Asis-Cruz, J., & Limperopoulos, C. (2024). Separating group- and individual-level brain signatures in the newborn functional connectome: A deep learning approach. NeuroImage, 299, 120806. 10.1016/j.neuroimage.2024.120806

Kim, J.-H., Zhang, Y., Han, K., Wen, Z., Choi, M., & Liu, Z. (2021). Representation learning of resting state fMRI with variational autoencoder. NeuroImage, 241, 118423. 10.1016/j.neuroimage.2021.118423

Kingma, D. P., & Ba, J. (2014). Adam: A method for stochastic optimization. arXiv Preprint arXiv:1412.6980.

Kingma, D. P., & Welling, M. (2013). Auto-Encoding Variational Bayes. CoRR. https://www.semanticscholar.org/paper/Auto-Encoding-Variational-Bayes-Kingma-Welling/5f5dc5b9a2ba710937e2c413b37b053cd673df02

Kong, R., Li, J., Orban, C., Sabuncu, M. R., Liu, H., Schaefer, A., Sun, N., Zuo, X.-N., Holmes, A. J., Eickhoff, S. B., & Yeo, B. T. T. (2019). Spatial Topography of Individual-Specific Cortical Networks Predicts Human Cognition, Personality, and Emotion. Cerebral Cortex, 29(6), 2533–2551. 10.1093/cercor/bhy123

Kong, R., Yang, Q., Gordon, E., Xue, A., Yan, X., Orban, C., Zuo, X.-N., Spreng, N., Ge, T., Holmes, A., Eickhoff, S., & Yeo, B. T. T. (2021). Individual-Specific Areal-Level Parcellations Improve Functional Connectivity Prediction of Behavior. Cerebral Cortex, 31(10), 4477–4500. 10.1093/cercor/bhab101

Kraus, B. T., Perez, D., Ladwig, Z., Seitzman, B. A., Dworetsky, A., Petersen, S. E., & Gratton, C. (2021). Network variants are similar between task and rest states. NeuroImage, 229, 117743. 10.1016/j.neuroimage.2021.117743

Kriegeskorte, N., Mur, M., & Bandettini, P. A. (2008). Representational similarity analysis—Connecting the branches of systems neuroscience. Frontiers in Systems Neuroscience, 2. 10.3389/neuro.06.004.2008

Kuhn, H. W. (1955). The Hungarian method for the assignment problem. Naval Research Logistics Quarterly, 2(1–2), 83–97. 10.1002/nav.3800020109

Labonte, A. K., Camacho, M. C., Moser, J., Koirala, S., Laumann, T. O., Marek, S., Fair, D., & Sylvester, C. M. (2024). Precision Functional Mapping to Advance Developmental Psychiatry Research. Biological Psychiatry Global Open Science, 4(6), 100370. 10.1016/j.bpsgos.2024.100370

Lancaster, J. L., Glass, T. G., Lankipalli, B. R., Downs, H., Mayberg, H., & Fox, P. T. (1995). A modality-independent approach to spatial normalization of tomographic images of the human brain. Human Brain Mapping, 3(3), 209–223. 10.1002/hbm.460030305

Langs, G., Golland, P., Tie, Y., Rigolo, L., & Golby, A. J. (2010). Functional Geometry Alignment and Localization of Brain Areas. Advances in Neural Information Processing Systems, 1, 1225–1233. https://www.ncbi.nlm.nih.gov/pmc/articles/PMC4010233/

Langs, G., Wang, D., Golland, P., Mueller, S., Pan, R., Sabuncu, M. R., Sun, W., Li, K., & Liu, H. (2016). Identifying Shared Brain Networks in Individuals by Decoupling Functional and Anatomical Variability. Cerebral Cortex, 26(10), 4004–4014. 10.1093/cercor/bhv189

Larivière, S., Vos de Wael, R., Hong, S.-J., Paquola, C., Tavakol, S., Lowe, A. J., Schrader, D. V., & Bernhardt, B. C. (2020). Multiscale Structure–Function Gradients in the Neonatal Connectome. Cerebral Cortex, 30(1), 47–58. 10.1093/cercor/bhz069

Laumann, T. O., Gordon, E. M., Adeyemo, B., Snyder, A. Z., Joo, S. J., Chen, M.-Y., Gilmore, A. W., McDermott, K. B., Nelson, S. M., Dosenbach, N. U. F., Schlaggar, B. L., Mumford, J. A., Poldrack, R. A., & Petersen, S. E. (2015). Functional System and Areal Organization of a Highly Sampled Individual Human Brain. Neuron, 87(3), 657–670. 10.1016/j.neuron.2015.06.037

Li, H., Satterthwaite, T. D., & Fan, Y. (2017). Large-scale sparse functional networks from resting state fMRI. NeuroImage, 156, 1–13. 10.1016/j.neuroimage.2017.05.004

Li, M., Wang, D., Ren, J., Langs, G., Stoecklein, S., Brennan, B. P., Lu, J., Chen, H., & Liu, H. (2019). Performing group-level functional image analyses based on homologous functional regions mapped in individuals. PLOS Biology, 17(3), e2007032. 10.1371/journal.pbio.2007032

Lynch, C. J., Elbau, I. G., Ng, T., Ayaz, A., Zhu, S., Wolk, D., Manfredi, N., Johnson, M., Chang, M., Chou, J., Summerville, I., Ho, C., Lueckel, M., Bukhari, H., Buchanan, D., Victoria, L. W., Solomonov, N., Goldwaser, E., Moia, S., … Liston, C. (2024). Frontostriatal salience network expansion in individuals in depression. Nature, 633(8030), 624–633. 10.1038/s41586-024-07805-2

Lynch, C. J., Elbau, I. G., Ng, T. H., Wolk, D., Zhu, S., Ayaz, A., Power, J. D., Zebley, B., Gunning, F. M., & Liston, C. (2022). Automated optimization of TMS coil placement for personalized functional network engagement. Neuron, 110(20), 3263–3277.e4. 10.1016/j.neuron.2022.08.012

Makhzani, A., Shlens, J., Jaitly, N., Goodfellow, I., & Frey, B. (2016, May 25). Adversarial Autoencoders. 10.48550/arXiv.1511.05644

Margulies, D. S., Ghosh, S. S., Goulas, A., Falkiewicz, M., Huntenburg, J. M., Langs, G., Bezgin, G., Eickhoff, S. B., Castellanos, F. X., Petrides, M., Jefferies, E., & Smallwood, J. (2016). Situating the default-mode network along a principal gradient of macroscale cortical organization. Proceedings of the National Academy of Sciences, 113(44), 12574–12579. 10.1073/pnas.1608282113

Miezin, F. M., Maccotta, L., Ollinger, J. M., Petersen, S. E., & Buckner, R. L. (2000). Characterizing the Hemodynamic Response: Effects of Presentation Rate, Sampling Procedure, and the Possibility of Ordering Brain Activity Based on Relative Timing. NeuroImage, 11(6), 735–759. 10.1006/nimg.2000.0568

Mitra, A., Snyder, A. Z., Tagliazucchi, E., Laufs, H., Elison, J., Emerson, R. W., Shen, M. D., Wolff, J. J., Botteron, K. N., Dager, S., Estes, A. M., Evans, A., Gerig, G., Hazlett, H. C., Paterson, S. J., Schultz, R. T., Styner, M. A., Zwaigenbaum, L., Network, T. I., … Raichle, M. (2017). Resting-state fMRI in sleeping infants more closely resembles adult sleep than adult wakefulness. PLOS ONE, 12(11), e0188122. 10.1371/journal.pone.0188122

Moore, L. A., Hermosillo, R. J. M., Feczko, E., Moser, J., Koirala, S., Allen, M. C., Buss, C., Conan, G., Juliano, A. C., Marr, M., Miranda-Dominguez, O., Mooney, M., Myers, M., Rasmussen, J., Rogers, C. E., Smyser, C. D., Snider, K., Sylvester, C., Thomas, E., … Graham, A. M. (2024). Towards personalized precision functional mapping in infancy. Imaging Neuroscience, 2, 1–20. 10.1162/imag_a_00165

Mueller, S., Wang, D., Fox, M. D., Yeo, B. T. T., Sepulcre, J., Sabuncu, M. R., Shafee, R., Lu, J., & Liu, H. (2013). Individual Variability in Functional Connectivity Architecture of the Human Brain. Neuron, 77(3), 586–595. 10.1016/j.neuron.2012.12.028

Muldoon, S. F., & Bassett, D. S. (2016). Network and Multilayer Network Approaches to Understanding Human Brain Dynamics. Philosophy of Science, 83(5), 710–720. 10.1086/687857

Myers, M. J., Labonte, A. K., Gordon, E. M., Laumann, T. O., Tu, J. C., Wheelock, M. D., Nielsen, A. N., Schwarzlose, R. F., Camacho, M. C., Alexopoulos, D., Warner, B. B., Raghuraman, N., Luby, J. L., Barch, D. M., Fair, D. A., Petersen, S. E., Rogers, C. E., Smyser, C. D., & Sylvester, C. M. (2024). Functional parcellation of the neonatal cortical surface. Cerebral Cortex, 34(2), bhae047. 10.1093/cercor/bhae047

Nair, V., & Hinton, G. E. (2010). Rectified linear units improve restricted boltzmann machines. 807–814.

Nenning, K.-H., Xu, T., Schwartz, E., Arroyo, J., Woehrer, A., Franco, A. R., Vogelstein, J. T., Margulies, D. S., Liu, H., Smallwood, J., Milham, M. P., & Langs, G. (2020). Joint embedding: A scalable alignment to compare individuals in a connectivity space. NeuroImage, 222, 117232. 10.1016/j.neuroimage.2020.117232

Nguyen, T. T., Qian, X., Ng, E. K. K., Ong, M. Q. W., Ngoh, Z. M., Yeo, S. S. P., Lau, J. M., Tan, A. P., Broekman, B. F. P., Law, E. C., Gluckman, P. D., Chong, Y.-S., Cortese, S., Meaney, M. J., & Zhou, J. H. (2023). Variations in Cortical Functional Gradients Relate to Dimensions of Psychopathology in Preschool Children. Journal of the American Academy of Child & Adolescent Psychiatry, 0(0). 10.1016/j.jaac.2023.05.029

Pandarinath, C., O’Shea, D. J., Collins, J., Jozefowicz, R., Stavisky, S. D., Kao, J. C., Trautmann, E. M., Kaufman, M. T., Ryu, S. I., Hochberg, L. R., Henderson, J. M., Shenoy, K. V., Abbott, L. F., & Sussillo, D. (2018). Inferring single-trial neural population dynamics using sequential auto-encoders. Nature Methods, 15(10), 805–815. 10.1038/s41592-018-0109-9

Pang, J. C., Aquino, K. M., Oldehinkel, M., Robinson, P. A., Fulcher, B. D., Breakspear, M., & Fornito, A. (2023). Geometric constraints on human brain function. Nature, 618(7965), Article 7965. 10.1038/s41586-023-06098-1

Petersen, S. E., Seitzman, B. A., Nelson, S. M., Wig, G. S., & Gordon, E. M. (2024). Principles of cortical areas and their implications for neuroimaging. Neuron, 0(0). 10.1016/j.neuron.2024.05.008

Poldrack, R. A., Laumann, T. O., Koyejo, O., Gregory, B., Hover, A., Chen, M.-Y., Gorgolewski, K. J., Luci, J., Joo, S. J., Boyd, R. L., Hunicke-Smith, S., Simpson, Z. B., Caven, T., Sochat, V., Shine, J. M., Gordon, E., Snyder, A. Z., Adeyemo, B., Petersen, S. E., … Mumford, J. A. (2015). Long-term neural and physiological phenotyping of a single human. Nature Communications, 6(1), 8885. 10.1038/ncomms9885

Pospelov, N., Tetereva, A., Martynova, O., & Anokhin, K. (2021). The Laplacian eigenmaps dimensionality reduction of fMRI data for discovering stimulus-induced changes in the resting-state brain activity. Neuroimage: Reports, 1(3), 100035. 10.1016/j.ynirp.2021.100035

Power, J. D., Barnes, K. A., Snyder, A. Z., Schlaggar, B. L., & Petersen, S. E. (2012). Spurious but systematic correlations in functional connectivity MRI networks arise from subject motion. NeuroImage, 59(3), 2142–2154. 10.1016/j.neuroimage.2011.10.018

Power, J. D., Cohen, A. L., Nelson, S. M., Wig, G. S., Barnes, K. A., Church, J. A., Vogel, A. C., Laumann, T. O., Miezin, F. M., Schlaggar, B. L., & Petersen, S. E. (2011). Functional Network Organization of the Human Brain. Neuron, 72(4), 665–678. 10.1016/j.neuron.2011.09.006

Power, J. D., Mitra, A., Laumann, T. O., Snyder, A. Z., Schlaggar, B. L., & Petersen, S. E. (2014). Methods to detect, characterize, and remove motion artifact in resting state fMRI. NeuroImage, 84, 320–341. 10.1016/j.neuroimage.2013.08.048

Puxeddu, M. G., Faskowitz, J., Betzel, R. F., Petti, M., Astolfi, L., & Sporns, O. (2020). The modular organization of brain cortical connectivity across the human lifespan. NeuroImage, 218, 116974. 10.1016/j.neuroimage.2020.116974

Raichle, M. E. (2015). The Brain’s Default Mode Network. Annual Review of Neuroscience, 38(1), 433–447. 10.1146/annurev-neuro-071013-014030

Ross, D. A., Lim, J., Lin, R.-S., & Yang, M.-H. (2008). Incremental Learning for Robust Visual Tracking. International Journal of Computer Vision, 77(1–3), 125–141. 10.1007/s11263-007-0075-7

Rousseeuw, P. J. (1987). Silhouettes: A graphical aid to the interpretation and validation of cluster analysis. Journal of Computational and Applied Mathematics, 20, 53–65. 10.1016/0377-0427(87)90125-7

Schaefer, A., Kong, R., Gordon, E. M., Laumann, T. O., Zuo, X.-N., Holmes, A. J., Eickhoff, S. B., & Yeo, B. T. T. (2018). Local-Global Parcellation of the Human Cerebral Cortex from Intrinsic Functional Connectivity MRI. Cerebral Cortex (New York, NY), 28(9), 3095–3114. 10.1093/cercor/bhx179

Scheinost, D., Kwon, S. H., Shen, X., Lacadie, C., Schneider, K. C., Dai, F., Ment, L. R., & Constable, R. T. (2016). Preterm birth alters neonatal, functional rich club organization. Brain Structure and Function, 221(6), 3211–3222. 10.1007/s00429-015-1096-6

Seitzman, B. A., Gratton, C., Laumann, T. O., Gordon, E. M., Adeyemo, B., Dworetsky, A., Kraus, B. T., Gilmore, A. W., Berg, J. J., Ortega, M., Nguyen, A., Greene, D. J., McDermott, K. B., Nelson, S. M., Lessov-Schlaggar, C. N., Schlaggar, B. L., Dosenbach, N. U. F., & Petersen, S. E. (2019). Trait-like variants in human functional brain networks. Proceedings of the National Academy of Sciences, 116(45), 22851–22861. 10.1073/pnas.1902932116

Seitzman, B. A., Snyder, A. Z., Leuthardt, E. C., & Shimony, J. S. (2019). The State of Resting State Networks. Topics in Magnetic Resonance Imaging : TMRI, 28(4), 189–196. 10.1097/RMR.0000000000000214

Shen, X., Tokoglu, F., Papademetris, X., & Constable, R. T. (2013). Groupwise whole-brain parcellation from resting-state fMRI data for network node identification. NeuroImage, 82, 403–415. 10.1016/j.neuroimage.2013.05.081

Shi, F., Salzwedel, A. P., Lin, W., Gilmore, J. H., & Gao, W. (2018). Functional Brain Parcellations of the Infant Brain and the Associated Developmental Trends. Cerebral Cortex, 28(4), 1358–1368. 10.1093/cercor/bhx062

Smith, S. M., Jenkinson, M., Woolrich, M. W., Beckmann, C. F., Behrens, T. E. J., Johansen-Berg, H., Bannister, P. R., De Luca, M., Drobnjak, I., Flitney, D. E., Niazy, R. K., Saunders, J., Vickers, J., Zhang, Y., De Stefano, N., Brady, J. M., & Matthews, P. M. (2004). Advances in functional and structural MR image analysis and implementation as FSL. NeuroImage, 23, S208–S219. 10.1016/j.neuroimage.2004.07.051

Stringer, C., Pachitariu, M., Steinmetz, N., Reddy, C. B., Carandini, M., & Harris, K. D. (2019). Spontaneous behaviors drive multidimensional, brainwide activity. Science, 364(6437), eaav7893. 10.1126/science.aav7893

Sun, L., Zhao, T., Liang, X., Xia, M., Li, Q., Liao, X., Gong, G., Wang, Q., Pang, C., Yu, Q., Bi, Y., Chen, P., Chen, R., Chen, Y., Chen, T., Cheng, J., Cheng, Y., Cui, Z., Dai, Z., … He, Y. (2023). Functional connectome through the human life span (p. 2023.09.12.557193). bioRxiv. 10.1101/2023.09.12.557193

Sydnor, V. J., Larsen, B., Bassett, D. S., Alexander-Bloch, A., Fair, D. A., Liston, C., Mackey, A. P., Milham, M. P., Pines, A., Roalf, D. R., Seidlitz, J., Xu, T., Raznahan, A., & Satterthwaite, T. D. (2021). Neurodevelopment of the association cortices: Patterns, mechanisms, and implications for psychopathology. Neuron, 109(18), 2820–2846. 10.1016/j.neuron.2021.06.016

Sylvester, C. M., Kaplan, S., Myers, M. J., Gordon, E. M., Schwarzlose, R. F., Alexopoulos, D., Nielsen, A. N., Kenley, J. K., Meyer, D., Yu, Q., Graham, A. M., Fair, D. A., Warner, B. B., Barch, D. M., Rogers, C. E., Luby, J. L., Petersen, S. E., & Smyser, C. D. (2022). Network-specific selectivity of functional connections in the neonatal brain. Cerebral Cortex, bhac202. 10.1093/cercor/bhac202

Tagliazucchi, E., von Wegner, F., Morzelewski, A., Brodbeck, V., Jahnke, K., & Laufs, H. (2013). Breakdown of long-range temporal dependence in default mode and attention networks during deep sleep. Proceedings of the National Academy of Sciences, 110(38), 15419–15424. 10.1073/pnas.1312848110

Talairach, J., & Tournoux, P. (1988). Co-planar Stereotaxic Atlas of the Human Brain: 3-dimensional Proportional System : an Approach to Cerebral Imaging. G. Thieme.

Tian, Y., Margulies, D. S., Breakspear, M., & Zalesky, A. (2020). Topographic organization of the human subcortex unveiled with functional connectivity gradients. Nature Neuroscience, 23(11), Article 11. 10.1038/s41593-020-00711-6

Tu, J. C., Myers, M., Li, W., Li, J., Wang, X., Dierker, D., Day, T., Snyder, A. Z., Latham, A., Kenley, J. K., & others. (2024). Early life neuroimaging: The generalizability of cortical area parcellations across development. bioRxiv : The Preprint Server for Biology, 2024–09. 10.1101/2024.09.09.612056

Tu, J. C., Wang, Y., Wang, X., Dierker, D., Sobolewski, C. M., Day, T. K. M., Kardan, O., Miranda-Domínguez, Ó., Moore, L. A., Feczko, E., Fair, D. A., Elison, J. T., Gordon, E. M., Laumann, T. O., Eggebrecht, A. T., & Wheelock, M. D. (2024). A Subset of Cortical Areas Exhibit Adult-like Functional Network Patterns in Early Childhood (p. 2024.07.31.606025). bioRxiv. 10.1101/2024.07.31.606025

Uddin, L. Q., Clare Kelly, A. m., Biswal, B. B., Xavier Castellanos, F., & Milham, M. P. (2009). Functional connectivity of default mode network components: Correlation, anticorrelation, and causality. Human Brain Mapping, 30(2), 625–637. 10.1002/hbm.20531

Van Essen, D. C., Drury, H. A., Dickson, J., Harwell, J., Hanlon, D., & Anderson, C. H. (2001). An Integrated Software Suite for Surface-based Analyses of Cerebral Cortex. Journal of the American Medical Informatics Association, 8(5), 443–459. 10.1136/jamia.2001.0080443

Van Essen, D. C., Glasser, M. F., Dierker, D. L., Harwell, J., & Coalson, T. (2012). Parcellations and Hemispheric Asymmetries of Human Cerebral Cortex Analyzed on Surface-Based Atlases. Cerebral Cortex, 22(10), 2241–2262. 10.1093/cercor/bhr291

Van Essen, D. C., Ugurbil, K., Auerbach, E., Barch, D., Behrens, T. E. J., Bucholz, R., Chang, A., Chen, L., Corbetta, M., Curtiss, S. W., Della Penna, S., Feinberg, D., Glasser, M. F., Harel, N., Heath, A. C., Larson-Prior, L., Marcus, D., Michalareas, G., Moeller, S., … Yacoub, E. (2012). The Human Connectome Project: A data acquisition perspective. NeuroImage, 62(4), 2222–2231. 10.1016/j.neuroimage.2012.02.018

Vanderwal, T., Eilbott, J., Finn, E. S., Craddock, R. C., Turnbull, A., & Castellanos, F. X. (2017). Individual differences in functional connectivity during naturalistic viewing conditions. NeuroImage, 157, 521–530. 10.1016/j.neuroimage.2017.06.027

Vos de Wael, R., Benkarim, O., Paquola, C., Lariviere, S., Royer, J., Tavakol, S., Xu, T., Hong, S.-J., Langs, G., Valk, S., Misic, B., Milham, M., Margulies, D., Smallwood, J., & Bernhardt, B. C. (2020). BrainSpace: A toolbox for the analysis of macroscale gradients in neuroimaging and connectomics datasets. Communications Biology, 3(1), Article 1. 10.1038/s42003-020-0794-7

Wang, D., Buckner, R. L., Fox, M. D., Holt, D. J., Holmes, A. J., Stoecklein, S., Langs, G., Pan, R., Qian, T., Li, K., Baker, J. T., Stufflebeam, S. M., Wang, K., Wang, X., Hong, B., & Liu, H. (2015). Parcellating cortical functional networks in individuals. Nature Neuroscience, 18(12), Article 12. 10.1038/nn.4164

Wang, F., Zhang, H., Wu, Z., Hu, D., Zhou, Z., Girault, J. B., Wang, L., Lin, W., & Li, G. (2023). Fine-grained functional parcellation maps of the infant cerebral cortex. eLife, 12, e75401. 10.7554/eLife.75401

Wen, X., Yang, M., Qi, S., Wu, X., & Zhang, D. (2024). Automated individual cortical parcellation via consensus graph representation learning. NeuroImage, 293, 120616. 10.1016/j.neuroimage.2024.120616

Wig, G. S. (2017). Segregated Systems of Human Brain Networks. Trends in Cognitive Sciences, 21(12), 981–996. 10.1016/j.tics.2017.09.006

Xia, Y., Xia, M., Liu, J., Liao, X., Lei, T., Liang, X., Zhao, T., Shi, Z., Sun, L., Chen, X., Men, W., Wang, Y., Pan, Z., Luo, J., Peng, S., Chen, M., Hao, L., Tan, S., Gao, J.-H., … He, Y. (2022). Development of functional connectome gradients during childhood and adolescence. Science Bulletin, 67(10), 1049–1061. 10.1016/j.scib.2022.01.002

Yeo, B. T. T., Krienen, F. M., Chee, M. W. L., & Buckner, R. L. (2014). Estimates of segregation and overlap of functional connectivity networks in the human cerebral cortex. NeuroImage, 88, 212–227. 10.1016/j.neuroimage.2013.10.046

Yeo, B. T. T., Krienen, F. M., Sepulcre, J., Sabuncu, M. R., Lashkari, D., Hollinshead, M., Roffman, J. L., Smoller, J. W., Zöllei, L., Polimeni, J. R., Fischl, B., Liu, H., & Buckner, R. L. (2011). The organization of the human cerebral cortex estimated by intrinsic functional connectivity. Journal of Neurophysiology, 106(3), 1125–1165. 10.1152/jn.00338.2011

Zeisel, A., Muñoz-Manchado, A. B., Codeluppi, S., Lönnerberg, P., La Manno, G., Juréus, A., Marques, S., Munguba, H., He, L., Betsholtz, C., Rolny, C., Castelo-Branco, G., Hjerling-Leffler, J., & Linnarsson, S. (2015). Cell types in the mouse cortex and hippocampus revealed by single-cell RNA-seq. Science, 347(6226), 1138–1142. 10.1126/science.aaa1934

Zhakubayev, A., & Hamerly, G. (2022). Clustering faster and better with projected data. 1–6.

